# Unbiased metabolic flux inference through combined thermodynamic and ^13^C flux analysis

**DOI:** 10.1101/2020.06.29.177063

**Authors:** Joana Saldida, Anna Paola Muntoni, Daniele de Martino, Georg Hubmann, Bastian Niebel, A. Mareike Schmidt, Alfredo Braunstein, Andreas Milias-Argeitis, Matthias Heinemann

## Abstract

Quantification of cellular metabolic fluxes, for instance with ^13^C-metabolic flux analysis, is highly important for applied and fundamental metabolic research. A current challenge in ^13^C-flux analysis is that the available experimental data are usually insufficient to resolve metabolic fluxes in large metabolic networks without making assumptions on flux directions and reversibility. To infer metabolic fluxes in a more unbiased manner, we devised an approach that does not require such assumptions. The developed three-step approach integrates thermodynamics, metabolome, physiological data, and ^13^C labelling data, and involves a novel method to comprehensively sample the complex thermodynamically-constrained metabolic flux space. Applying our approach to budding yeast with its compartmentalised metabolism and parallel pathways, we could resolve metabolic fluxes in an unbiased manner, we obtained an uncertainty estimate for each flux, and we found novel flux patterns that until now had remained unknown, likely due to assumptions made in previous ^13^C flux analysis studies. We expect that our approach will be an important step forward to determine metabolic fluxes with improved accuracy in microorganisms and possibly also in more complex organisms.

## INTRODUCTION

Intracellular metabolic fluxes represent a major phenotypic output of many cellular processes (Sauer, 2006). Knowledge of metabolic fluxes is therefore highly important for fundamental metabolic research (Hackett et al, 2016;Hui et al, 2017), metabolic engineering, and biotechnology (Nielsen & Keasling, 2016) as well as for several areas of biomedicine (DeBerardinis & Chandel, 2016). The current gold standard to determine intracellular fluxes is ^13^C metabolic flux analysis (Antoniewicz, M. R., 2015;Buescher et al, 2015;Dalman et al, 2016;Long & Antoniewicz, 2019), which is commonly applied for microorganisms (Dahlin et al, 2019;Diaz et al, 2019;Ghosh et al, 2016;Haverkorn van Rijsewijk et al, 2011;Kohlstedt & Wittmann, 2019;Wasylenko & Stephanopoulos, 2015). To obtain intracellular fluxes with ^13^C metabolic flux analysis, a mass and isotopomer balance model is fitted to isotopomer abundance measurements obtained from ^13^C labelling experiments, as well as to physiological data, such as rates of growth, metabolite uptake and excretion (Nöh et al, 2008;Rantanen, 2008;Wiechert, W., 2001). The high number of isotopomer balances and their non-linearity make this method computationally intensive (Antoniewicz, 2015). As ^13^C labelling in metabolites can also ‘travel’ against the net flux direction of a reaction (through a reaction’s reversibility) (Wiechert, 2007), isotopomer models that are used to fit the labelling data also contain variables for these so-called ‘labelling exchange’ fluxes, adding additional degrees of freedom to the already challenging optimisation problem.

While ^13^C metabolic flux analysis is also applied at genome-scale (García Martí-n et al, 2015;Gopalakrishnan & Maranas, 2015), it turns out that the information contained in the measured ^13^C labelling patterns is relatively limited (Foguet et al, 2019). To deal with this fact, the degrees of freedom present in the used models are typically reduced, for instance by either using metabolic network models of small size or by making heuristic assumptions on reaction directions and reaction reversibility (i.e. on the labelling exchange fluxes). However, the basis of such assumptions is often unclear (Zamboni, 2011), and reaction directions and labelling exchange fluxes can also depend on the environmental conditions and on the organism under study. It has been stated that there is a significant risk in making erroneous assumptions (Martínez et al, 2014), which calls for approaches to tackle this issue (Theorell & Noh, 2020) or for an approach that does not require any such assumptions. The need for assumptions that constrain the degrees of freedom is even more pressing in eukaryotic organisms (Antoniewicz, 2015;Lehnen et al, 2017;Zamboni, 2011), where key metabolic pathways are distributed over the cytoplasm and mitochondria, where several reactions exist in both compartments and many metabolites are exchanged between the two compartments. Another important challenge in ^13^C flux analysis is also the quantification of the uncertainty in the estimated fluxes (Theorell et al, 2017).

A way to address these challenges in current ^13^C flux analysis could be to use the second law of thermodynamics together with metabolite concentration measurements to constrain reaction directions. Ideally, one would simultaneously fit physiological, metabolome, and ^13^C labelling data into a combined stoichiometric-thermodynamic-isotopomer model. However, we consider such integration highly challenging. The addition of the second law renders such model non-linear and the solution of mass and isotopomer models is already highly computationally demanding due to the large numbers of equations.

Here, to yet exploit thermodynamics and metabolome data for the estimation of metabolic fluxes through a realistically sized eukaryotic metabolic network without using any *a priori* assumptions, we developed an approach to integrate thermodynamics, metabolome and physiological data, and ^13^C labelling data. Specifically, we developed a three-step approach, including non-linear fittings with a thermodynamic and stoichiometric model (Niebel et al, 2019), a novel sampling approach, and fitting of an isotopomer model to ^13^C labelling data. With this approach, we were able to estimate metabolic fluxes and uncertainty quantification in an unbiased manner, where we found novel flux patterns that until now had remained unknown. We expect that our new approach to resolve intracellular fluxes will be an important step forward to determine metabolic fluxes with improved accuracy and less bias in microorganisms, and possibly also in more complex organisms.

## RESULTS

### Overview method

To infer intracellular metabolic fluxes, our approach makes use of different types of steady-state experimental data and mathematical constraints, and involves the following three steps (Fig. 1): First, we fit a combined stoichiometric and thermodynamic model (Niebel et al, 2019) to physiological data (rates of growth, metabolite uptake and excretion), metabolome data and estimates of Gibbs standard energies of reaction (Δ_*r*_*G*^0^) obtained from the component contribution method (Noor et al, 2013). This fitting delivers Δ_*r*_*G*^0^ values with a common thermodynamic reference state. Using the estimated Δ_*r*_*G*^0^ and the experimental data together with the model, we then perform variability analysis to determine upper and lower bounds for the model variables, i.e. fluxes, metabolite concentrations, and Gibbs energies of reaction (Δ_*r*_*G*). Such variable bounds demark the boundaries of the solution space constrained by stoichiometry, thermodynamics, metabolome, and physiological data. Second, we sample net flux solutions from this solution space employing a novel approach based on Markov-Chain Monte Carlo. A net flux solution is a set of flux values for each reaction in the network, consistent with the experimental data and model constraints. Third, we use each sampled net flux solution to fit an isotopomer model to ^13^C labelling data. This fitting allows us to score the quality of each net flux solution based on the fitting residual. Thereby, we can further close in on the metabolic fluxes as they occur in the organism under the respective conditions.

**Figure 1:**
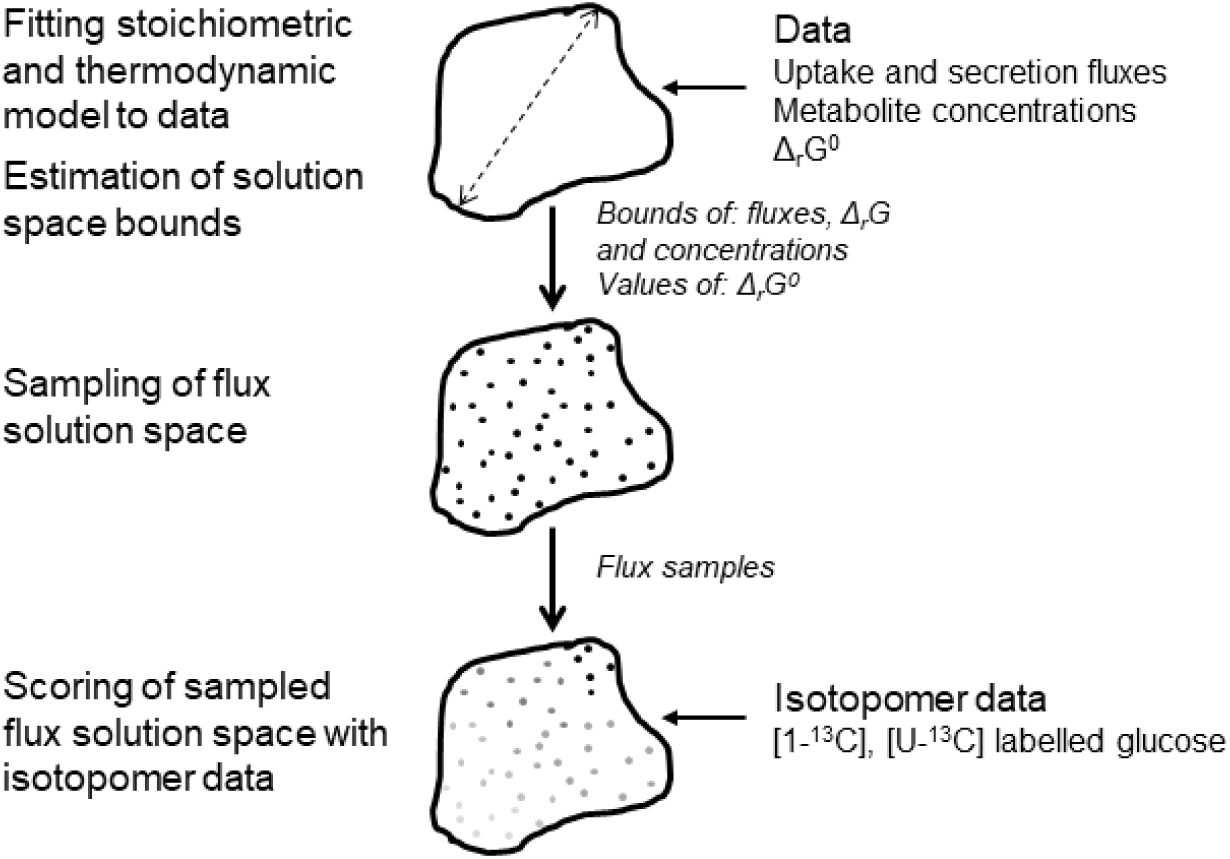
Illustration of our novel approach to estimate metabolic fluxes. First, a combined stoichiometric and thermodynamic metabolic model is fitted to metabolite uptake and secretion fluxes and metabolite concentrations to estimate a consistent set of Gibbs standard energies of reaction (Δ_*r*_*G*^0^), followed by determination of variable bounds – fluxes, concentrations and Gibbs energies of reaction (Δ_*r*_*G*) - through variability analysis with the Δ_*r*_*G*^0^ values fixed. Next, the solution space defined by the model equations and variable bounds is sampled with a novel method developed in this work to obtain samples of net fluxes. Finally, the net flux samples are scored based on the fitting of an isotopomer and mass balance model to isotopomer data from experiments with labelled glucose. The scores allow the identification of the most plausible flux distributions.

We developed the method using budding yeast *Saccharomyces cerevisiae* as an example for a eukaryotic metabolism. To demonstrate that the proposed method applies to different metabolic modes and to retrieve new biological insights from such comparison, we applied it to two yeast strains exhibiting either a fermentative (wildtype) and respiratory (TM6) metabolism. The latter strain uses respiration because it has a low rate of glucose uptake, as it expresses only a single chimeric hexose transporter (Elbing et al, 2004).

### Metabolome, physiological data, and thermodynamics to constrain the flux space

For the first step of our approach, we use a combined stoichiometric and thermodynamic model (TSM) for budding yeast (Niebel et al, 2019), to which some additional reactions for uptake and metabolism of alternative nutrients were added (Supplementary Data File 1). The stoichiometric part of the model contains 258 reactions (*N*), encompassing the reactions of central and amino acid metabolism distributed between the cytosol and mitochondria, transport processes (*METRxn*), and metabolite uptake and secretion reactions (i.e. so-called ‘exchange reactions’, *EXGRxn*). As the precise stoichiometry of transport processes is often unknown (Bianchi et al, 2019), our model includes several stoichiometric variants for the transport processes, e.g. variants with and without co- /anti-transported protons. Due to several reactions occurring in linear pathways, a flux solution in our model is defined by 148 independent reactions. 156 unique metabolites are balanced in different compartments, yielding 206 steady-state mass balance equations (*M*) defined by

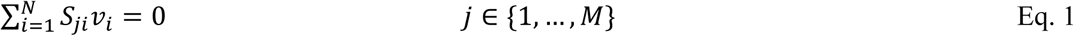

where *S* is the matrix containing the stoichiometric coefficients for metabolite *j* in reaction *i*, and *v* is the metabolic flux through reaction *i*. The model also includes pH and charge balances for each compartment. The charge and number of protons of each metabolite are defined by their pKa value and the pH in the respective compartment.

The thermodynamic part of the model consists of the second law of thermodynamics,

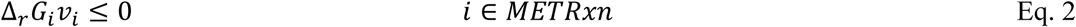

where Δ_*r*_*G* is the Gibbs energy of reaction *i*, made a function of metabolite concentrations through

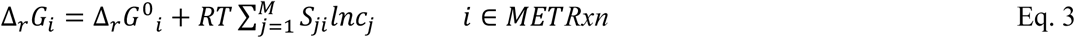

where Δ_*r*_*G*^0^ is the standard Gibbs energy of reaction, *R* the gas constant, *T* the temperature and *ln c* is the logarithm of the concentration of metabolite *j*, assuming a dilute system. The Gibbs energies of transport processes are modelled as done previously (Jol et al, 2010). Notably, in our model, all reactions and processes are considered bidirectional and the reaction directions are only constrained by the second law of thermodynamics (Eq. 2). The model also includes a Gibbs energy balance, which ensures that the Gibbs energy that is dissipated in the metabolic reactions and transport processes is transported to the environment, through

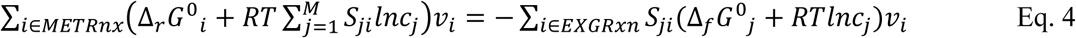

Where Δ_*f*_*G*^0^ is the standard Gibbs energy of formation of metabolite *j*. Variables of the model are fluxes (v), metabolite concentrations (c), Gibbs energies of reaction (Δ_*r*_*G*), and for the first fitting in Fig. 1, also standard Gibbs energies of reaction (Δ_*r*_*G*^0^).

We fitted this model to experimental data to obtain a complete and consistent set of standard Gibbs energies of reaction (Δ_*r*_*G*^0^) values and to define the limits of the solution space of fluxes, metabolite concentrations and Gibbs reaction energies (Δ_*r*_*G*) for each of the two metabolic modes (i.e. the wildtype and the TM6 mutant strain). For each strain, the experimental data consisted of metabolite concentrations measured for 31 metabolites, physiology data (growth rate, metabolite uptake, and secretion rates) (Supplementary Data File 2), as well as Δ_*r*_*G*^0^ values and corresponding estimation errors, as estimated from the Component Contribution Method (Noor et al, 2013), for 171 of the 178 Gibbs energy-dissipating reactions. To fit the model to the cell-averaged metabolite data, we included equations that combine the cytosolic and mitochondrial metabolite concentrations into cell-average metabolite concentrations, using known volume fractions between the cellular compartments (Niebel et al, 2019). In this fitting, we also included the loop law To ensure that the estimated Δ_*r*_*G*^0^ values were in the same reference state, we also enforced the loop law (Beard et al, 2002) as additional constraint for in the fitting, as done in (Niebel et al, 2019).

We fitted the model to the experimental data by minimising the sum of square differences between measured and predicted values, weighted by the corresponding standard deviations. This fitting was performed as previously (Niebel et al, 2019), except that here we did not use regularisation due to the limited number of unknown Δ_*r*_*G*^0^ values. We fitted the two experimental data sets (from WT and TM6) simultaneously, to enforce one joint set of Δ_*r*_*G*^0^ for both strains, while the rest of the model variables were allowed to be strain-dependent. We found that the model could fit well to both sets of data (Fig. 2a,b) and that the Δ_*r*_*G*^0^ values only marginally deviate from the values obtained from the component contribution method (Fig. 2c). For our following analyses, we fixed the Δ_*r*_*G*^0^ to the values estimated in this fitting and we kept Eq. 4 (see Supplementary Information).

**Figure 2:**
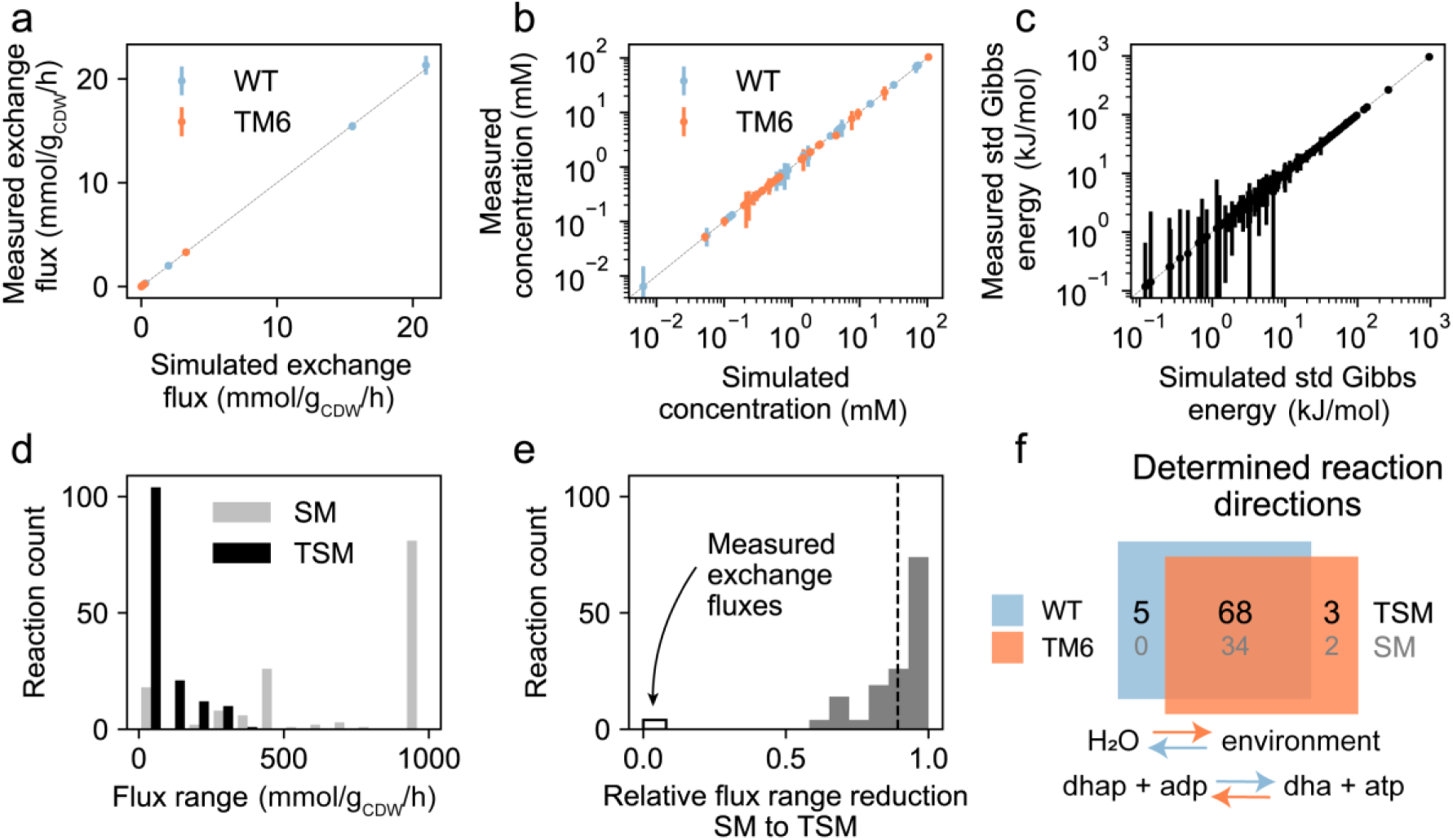
Metabolome, physiological data and thermodynamics constrain the flux solution space. (a), (b), (c) Measured against simulated (fitted) values for physiological data (exchange fluxes) (a), metabolite concentrations (b) and standard Gibbs reaction energies, Δ_*r*_*G*^0^ (c). Experimental data in (a) and (b) were measured in this work and data in (c) was calculated with the component contribution method (Noor et al, 2013). Colours represent the two strains (WT, TM6) and vertical bars represent experimental standard deviations calculated from three experiments for (a) and (b), and errors from Component Contribution Method for (c). (d) Histograms of flux ranges, constrained by a stoichiometric model (SM) and physiological data, and a combined thermodynamic/stoichiometric model (TSM) and metabolome and physiological data. Flux ranges were calculated as upper bound minus lower bound, where the bounds were determined from variability analyses (Mahadevan & Schilling, 2003). (e) Histogram of relative flux range reduction from SM to TSM, calculated as (range_SM_ - range_TSM_) / range_SM_. The dashed line indicates the mean value of the histogram excluding measured exchange fluxes, which are represented with the white bar on the left. (f) Diagram with the number of identified reaction directions for WT (in blue) and TM6 (in orange) where numbers in black and grey correspond to results obtained with TSM and SM, respectively. The two reactions shown at the bottom operate in opposite directions in WT and TM6: water transport to the environment and dihydroxyacetone kinase. The blue arrows indicate the direction as it occurs in the WT, and the orange TM6.

Next, to determine the bounds of the solution space of fluxes v, log-metabolite concentrations ln(c) and Δ_*r*_*G*, as defined by the above-introduced equations and data, we performed variability analysis (Mahadevan & Schilling, 2003) for WT and TM6, yielding upper and lower bounds for all variables (Supplementary Data File 3). For the variability analyses, we constrained the measured variables (i.e. exchange fluxes and metabolite concentrations) to the values estimated in the fitting plus/minus three experimental standard deviations. Subsequently, we maximised and minimised each variable in independent optimisations. The resulting variable bounds (Fig. S1a-c) define the limits of the strain-dependent solution spaces that are consistent with the experimental data and the strain-independent set of Δ_*r*_*G*^0^. The model fitting to the experimental data and the variability analysis were both performed in GAMS (General Algebraic Modeling System, Release 24.6).

To assess the size of the flux solution space of the wildtype, we calculated flux ranges (i.e. upper minus lower flux bound) and box volumes. The box volume is the volume of the hyperrectangle formed by the Cartesian product of the individual flux ranges, and was calculated by multiplying the lengths of all sides of this hyperrectangle. To investigate the extent to which the metabolite data and thermodynamics constrained the flux solution space, we also fitted solely the stoichiometric part of our model (i.e. Eq. 1, abbreviated as SM) to the physiological data, and similarly determined the flux bounds and the box volume. We found that the box volume of the flux solution space decreased from the SM to the TSM by a factor of 10^181^. This very large reduction can also be seen in the flux ranges, where flux ranges obtained from the SM were in general larger than the ones from the TSM (Fig. 2d) and their overall relative reduction was on average 89% (excluding the measured exchange fluxes; Fig. 2e). This huge reduction in flux ranges is due to the implemented thermodynamic constraints, which remove infeasible loops of reactions (De Martino, 2017). Significant flux range reductions occurred in all pathways (Fig. S1d).

The constraining effect of the metabolome data and thermodynamics can also be illustrated by reactions that are constrained in a particular direction under the two different metabolic modes. From the sign of the estimated flux bounds, obtained from the variability analyses, we identified the reactions constrained in a particular direction with the SM and the TSM. With the TSM, we found that, out of the 148 independent reactions, 96 WT and 94 TM6 reactions are constrained in one direction. While most pathways have similar fractions of identified reaction directions between the two strains, differences mostly occur in glycolysis and the pentose phosphate pathway (Fig. S1e). In contrast, with the SM, we could only determine directions (both positive and negative) of 34 (WT) and 36 (TM6) of the fluxes (Fig. 2f). Remarkably, two of the reaction directions determined with the TSM were opposite between WT and TM6: water transport and dihydroxyacetone kinase (Fig. 2f). In addition, eight (i.e. 5+3) reactions were determined to run in one direction under one metabolic mode, but they could still operate in both directions under the other.

These results show that exploiting metabolome data and thermodynamics significantly constrains the flux solution space, compared to when only physiological data and a stoichiometric model is used. Furthermore, although thermodynamics only influences flux directions, our analysis shows that the integration of thermodynamic constraints with metabolome and physiological data results in flux ranges being further constrained. Using thermodynamics and metabolome data also has the advantage that reaction directions do not need to be defined *ab initio*, which avoids bias in flux estimation.

### Assessment of the feasible solution space with a new sampling approach

The variable bounds that we had identified at this point could, in a next step, be used together with ^13^C labelling data to estimate fluxes through a mass and isotopomer model, i.e. for classical ^13^C MFA. However, given the large number of variables (i.e. net and labelling exchange fluxes) in the complex optimisation problem and the lack of constraints on labelling exchange fluxes and on net flux directions, the solution space was still too large for the optimiser to find a good fit. Thus, we needed to find an alternative approach. Specifically, we aimed to sample the net flux solution space, constrained with stoichiometry, thermodynamics, physiology, and metabolome data, and then to score such net flux samples in the next step with ^13^C labelling data. To this end, however, we first needed a suitable sampling method.

Our solution space is non-convex: while the mass balances (Eq. 1) and Δ_r_G as a function of log-metabolite concentrations (Eq. 3) are linear relationships, the second law of thermodynamics (Eq. 2) and the Gibbs energy balance (Eq. 4) are bilinear, making the solution space non-convex. Due to the bilinear terms, classical samplers based on Markov Chain Monte Carlo (MCMC) methods for sampling of flux solutions (Binns et al, 2015;Chaudhary et al, 2016;Price et al, 2006;Saa & Nielsen, 2016) cannot be used here. Moreover, an algorithm able to sample non-convex spaces, developed for a three-dimensional problem unrelated to metabolic networks (Zappa et al, 2018), was unsuitable for our problem.

Thus, we had to develop a sampling approach capable of sampling the high-dimensional non-convex solution space. To address this challenge, we took the following four steps (Fig. 3), which, in combination, allowed us to achieve our goal: (i) division of the non-convex space into two convex spaces (polytopes), (ii) division of the flux polytope into sectors, (iii) scaling of polytopes, and (iv) use of a parallel tempering Monte Carlo method. We will explain these measures in detail in the following.

**Figure 3:**
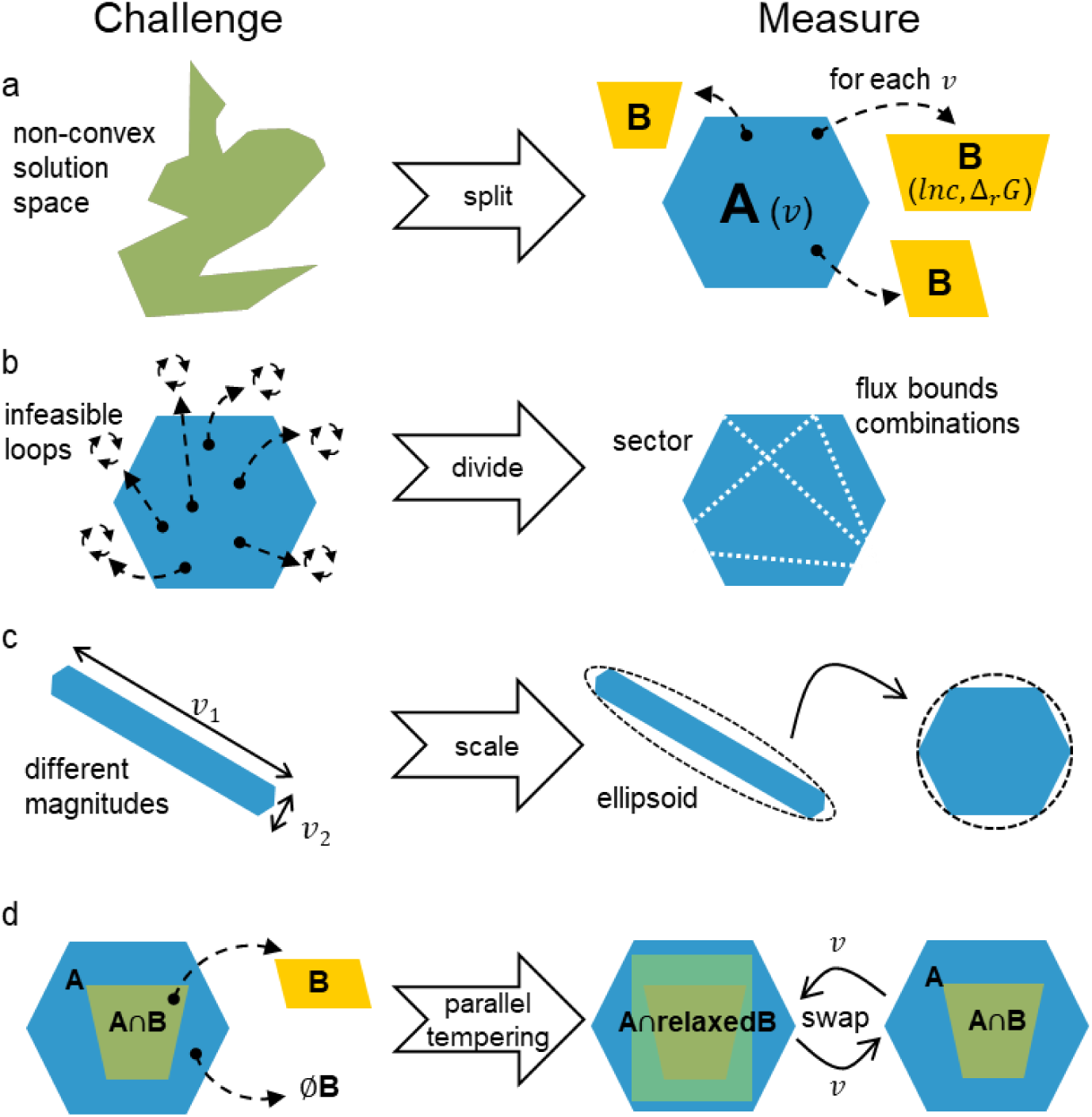
Overview of challenges and measures used to sample the thermodynamically constrained solution space. (a) The non-convexity of the solution space (in green) of fluxes, Gibbs energies and log-concentrations consistent with Eqs. 1-4 and respective bounds is overcome with a split of the space into two convex polytopes (A and B), based on the knowledge that for a given flux solution (sampled from flux polytope A, in blue) there is a polytope defined by thermodynamic constraints in the (ln c, Δ_*r*_*G*) solution space (polytope B, in yellow). (b) Thermodynamically infeasible reaction loops (represented by small three arrow circles) immensely decrease the acceptance rate of a hit-and-run sampler in Polytope A, which can be partly overcome through a division of the flux solution space in non-overlapping sectors that avoid most loops. The reactions involved in many infeasible loops were identified and such loops were removed by setting one of the flux bounds to 0. Sectors are then defined through combinations of flux bounds, illustrated by the white dashed lines. (c) Due to the diverse orders of magnitude of the fluxes (for example *v*_1_ is much larger than v_2_), it is challenging for a hit-and-run sampler to sample both fluxes to an equal extent. This issues can be resolved by scaling the flux solution space (A) with an ellipsoid transformation, where the ellipsoid enclosing the flux solution space is estimated (dashed line) and used to rescale the flux solution space, such that the different directions of the flux space have roughly equal magnitudes. We obtained the ellipsoid parameters from the covariance matrix of an approximate distribution of the metabolic fluxes, provided by the Expectation Propagation algorithm (Braunstein et al, 2017). (d) The size of the flux polytope A is much larger than the size of A∩B (flux polytope A conditional on existence of non-empty polytope B) causing the overall search for flux samples complying with B to be very inefficient. We use a parallel tempering cascade to increase the acceptance rate and the exploration of the sampler. The two green shapes with different shades represent replicas of the flux solution space with different levels of relaxation of constraints. By occasionally swapping flux solutions (v) between replicas, we make the exploration of the flux solution space more efficient.

First, to solve the issue with the non-convexity, we made use of the fact that Eqs. 2 and 4 have a bilinear structure, where if *v* is fixed the two equations become linear, making *ln c* and Δ_*r*_*G* lie inside a convex polytope. This fact allowed us to split the solution space into two convex parts, i.e. a flux polytope (A) and a concentration polytope (B) (Fig. 3a). We could then, first, sample the convex polytope (A) of fluxes (v) established by Eq. 1 and v bounds, and then for each v point in A sample the convex polytope (B) of concentrations (ln c) and Gibbs energies of reaction (Δ_*r*_*G*) generated by Eqs. 2, 3, 4 and ln c and Δ_*r*_*G* bounds.

The splitting of the sampling problem into two polytopes overcame non-convexity but raised the problem that the samples generated in the flux polytope contained many thermodynamically infeasible loops. Even though many reaction directions were already determined, the remaining reactions with in an undetermined direction could be involved in infeasible loops. To avoid sampling solutions with infeasible loops, we identified four major infeasibility-causing loops involving nine reactions, by solving the dual problem of the loop law equation iteratively (cf. Method section ‘Division of the flux solution space into sectors’). By imposing different combinations of flux bounds for these reactions (i.e. as either positive or negative), we divided the solution space into 16 non-overlapping sectors (Fig. 3b), i.e. subsets of the original flux space, each defined by a unique sign combination of the fluxes involved in infeasible loops. By sampling these sectors individually, we avoided sampling solutions that contained thermodynamically infeasible loops.

When sampling the flux solution space of each sector using a Hit-and-Run Markov Chain Monte Carlo (HR) algorithm (Chen & Schmeiser, 1996), we ran into the next problem, connected with the fact that the ranges of metabolic fluxes span several orders of magnitude (i.e. from 10^−5^ to 10^2^ mmol/g_DW_/h, Fig. S1a). This made it very difficult to sample the space uniformly in all dimensions (i.e. fluxes). To address this issue, we computed an ellipsoidal approximation of the feasible solution space of each sector by Expectation Propagation (Braunstein et al, 2017),, which we used to properly re-scale the space to facilitate sampling (Fig. 3c). Expectation Propagation provided a multivariate Gaussian approximation of the joint distribution of metabolic fluxes constrained by mass balance and flux bounds.

Despite this splitting of the flux solution space in sectors, when we sampled the flux polytope sectors with the HR algorithm and checked (with linear optimisation) whether a non-empty polytope B exists for the each flux sample, we found that only an exceedingly small fraction (0.001%) of the sampled flux configurations resulted in a non-empty polytope B. Consequently, sampling of polytope A via HR and accepting only the flux configurations with a non-empty polytope B, leads to a highly inefficient sampler. To increase the acceptance rate, we developed a Parallel Tempering Monte Carlo algorithm (Swendsen & Wang, 1986) (Fig. 3d). Using Parallel Tempering, we sampled simultaneously six replicas of the solution space (the subset of polytope A for which a non-empty polytope B exists), each with different relaxation degrees (‘temperatures’) for the constraints Eqs. 3 and 4. Replicas with higher temperature have higher constraint violation resulting in more extensive sampling of the solution space, while lower temperature replicas perform sampling closer to the target distribution (corresponding to the original constraints of Eqs. 3 and 4) and move more locally. During the sampling of the replicas, flux samples are swapped between solution spaces of adjacent relaxation degrees, according to the probability of constraint violation at adjacent temperatures (‘Metropolis move’). We manually adjusted the number of replicas and their temperatures to achieve an average swap rate of 20% among replicas at adjacent temperatures. Through this approach, we created a chain of samples passed down to less-relaxed solution spaces allowing for their efficient exploration. In this way, we could access regions of the solution space that we would not have been able to access with regular sampling approaches, and comprehensively explore the flux solution space much more efficiently than the simple rejection sampler described above.

Finally, to obtain the complete flux sample over the full flux solution space after sampling each sector individually, we combined all the sector samples using the volume of the respective sector as a weight. We calculate the sector volume using the approximation of the flux solution space determined by Expectation Propagation (Braunstein et al, 2017). This calculated volume reflects the number of feasible flux samples of a multivariate Gaussian distribution. Looking at the complete flux sample over the complete flux solution space, we found that three independent runs of the PT sampling approach gave highly similar results in terms of flux medians (Fig. 4a) and variability (Fig. 4b) across samples, and also for individual flux distributions (Fig. S2), which indicates that the sampling approach is reproducible.

**Figure 4:**
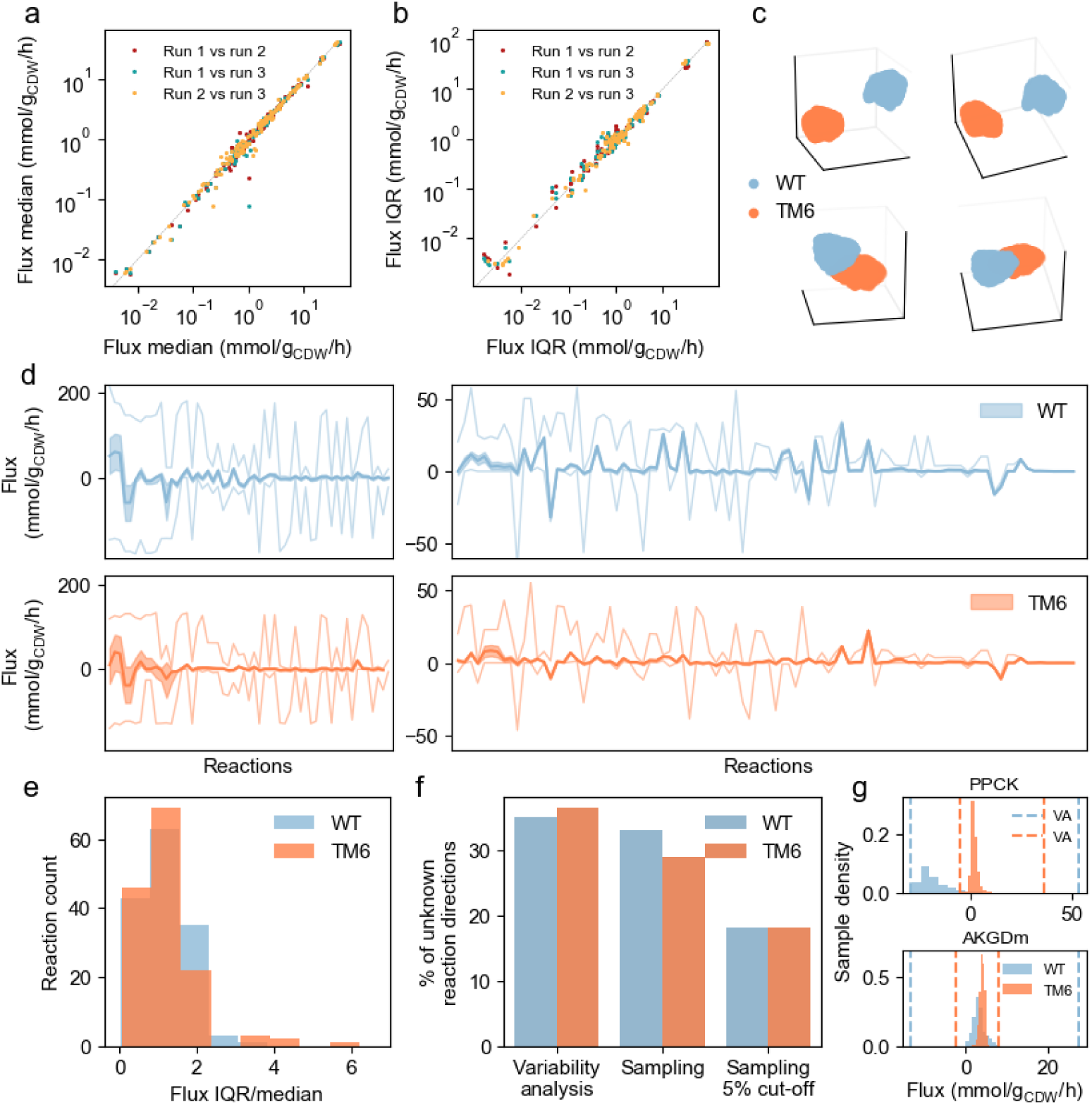
Sampling thermodynamically constrained flux solution space with the new sampling approach. (a), (b) Comparison of flux medians (a) and interquartile ranges (IQRs) (b) obtained for three independent runs of sampling, showing the sampler’s reproducibility (example for the sampling of TM6 flux solution space). Three comparisons (sets of points) between the three runs are presented in the plots in different colours. The diagonal dashed line represents the same value for the x and y axes. (c) Visualisation of the flux solution spaces of WT (in blue) and TM6 (in orange) from 4 different visualisation angles. Dimensionality reduction was performed with UMAP (Uniform Manifold Approximation and Projection for Dimension Reduction, (McInnes et al, 2018)), a technique which represents the points of a high-dimensional sample in a lower-dimensional space while preserving aspects of the global structure of the sample. (d) Flux data from sampling for all independent reactions for WT (top, in blue) and TM6 (bottom, in orange), respectively. Light colour lines represent flux bounds from variability analysis, coloured area represents the span of flux IQR and the dark line shows the median of the flux distribution. Reactions are arranged on the x-axis in decreasing order of magnitude of WT median fluxes. Left panels show reactions with high flux variability and right panels with low variability. (e) IQR/median of sampled fluxes for WT (in blue) and TM6 (in orange). Here, only IQR/median values up to 6 are shown; data with the more extended x-axis is shown in Fig. S3. (f) Percentage of undetermined flux directions for WT (in blue) and TM6 (in orange) after performing variability analysis (VA), after sampling, and after sampling and cutting off of 5% on the tail of the flux distributions, resulting in more determined flux directions. (g) Examples of flux distributions after the sampling and application of a 5% cut-off on the left and right distribution tails for both WT (in blue) and TM6 (in orange). Dashed lines represent variability (VA) flux bounds for WT and TM6 with the respective colours. PPCK: Phosphoenolpyruvate carboxykinase, AKGDm: mitochondrial oxoglutarate dehydrogenase. For these two reactions, the removal of the 5% tail constrained the reactions in one direction. For all plots, sampled fluxes were obtained from sampling the flux solution space conditional on the concentration space with Hit-and-Run and Parallel Tempering.

This new sampling approach allowed us to sample the complete flux solution space constrained by the thermodynamic and stoichiometric model and the physiological and metabolite data. It is important to note that with this approach we find flux samples that fulfil the thermodynamic constraints even though we are not simultaneously sampling concentrations and Gibbs energies, as we just check for the existence of a feasible solution in polytope B. As the sampling is performed with HR, the resulting flux samples are uniform in the flux solution space conditional on the feasibility of the concentration solution space.

With this sampling method, we generated 100 000 samples covering the complete flux solution space for each of the two strains (Supplementary Data File 4). While the two flux solution spaces are separated in a 3D reduction visualisation established by UMAP (McInnes et al, 2018) (Fig. 4c), a clear separation is not directly visible when the median and the interquartile range (IQR) of the marginal flux distributions is compared between the two strains (Fig. 4d). Yet, comparing the IQRs of the samples with the bounds determined by variability analysis, we find that the sampling confined the flux solution space much further (Fig. 4d). We also found that 71 and 96 of the fluxes of the independent reactions were well constrained, i.e. had an IQR/median lower than 1, for WT and TM6 respectively (Fig. 4e). When we compared the percentage of reactions with unknown reaction directions, we found that the percentage dropped from 35% and 36% from the variability analysis bounds to 33% and 29% after the sampling, for WT and TM6 respectively. When further cutting off the tail of the flux distributions by 5%, the latter percentages dropped to 18% (Fig. 4f). Finally, as exemplified with the fluxes through the phosphoenolpyruvate carboxykinase and the mitochondrial oxoglutarate dehydrogenase reaction, flux ranges are very much reduced compared to the bounds obtained from the variability analysis and they can be completely different between the two strains, as is the case for the phosphoenolpyruvate carboxykinase (Fig. 4g).

With our new sampling approach, we comprehensively assessed the flux solution space conditional on the concentration polytope. We also generated samples for metabolite concentrations and Δ_r_G for each flux sample (Supplementary Data File 4).

### Scoring net flux samples with ^13^C labelling data

Next, we aimed to score each flux sample against measured ^13^C labelling data, i.e. isotope abundances (Supplementary Data File 5, Supplementary Data File 6). Therefore, we implemented a mass and isotopomer balance model with a stoichiometry identical to the TSM model, but excluding those reactions that do not interconvert carbon atoms, and with only one stoichiometric variant for each transport process (Supplementary Data File 7). Importantly, this slightly reduced model retained identical carbon flow representation compared to the TSM model. For the isotopomer balances, an atom transition network was generated to simulate the carbon flow through the network. As ^13^C labelling in metabolites can also be exchanged against the net direction of a reaction (Wiechert, W. et al, 2001), the isotopomer model also included so-called ‘labelling exchange fluxes’ for each reaction (*v*^*xch*^), with *v*^*xch*^ = min(*v*^+^, *v*^−^), where (*v*^+^) and (*v*^−^) are the forward and backward flux of a reaction, respectively. The net flux (*v*^*net*^) is defined as *v*^*net*^ = *v*^+^ − *v*^−^. We implemented the mass and isotopomer model in FluxML format to use in the software 13CFlux2 (Weitzel et al, 2013).

Because there is no single optimal tracer experiment with enough information to determine all metabolic fluxes in complex networks (Antoniewicz, 2013), parallel labelling experiments with different labelled substrates are commonly performed to improve the resolution of fluxes in different pathways (Antoniewicz, 2015). Here, we measured amino acid labelling in an experiment where 100% of the glucose was labelled at the first carbon atom [1-^13^C], and in another experiment, where we used 80% unlabelled glucose and 20% fully labelled glucose [U-^13^C], as previously done (Zamboni et al, 2009). These two sets of labelling data could be used to fit the isotopomer model individually or simultaneously.

Next, we fitted the model to the ^13^C labelling data, where we used a net flux sample as input, leaving the labelling exchange fluxes as the sole free variables in the optimisation. The residual of the optimisation (i.e. the sum of square differences between measured and predicted abundances, weighted by their measurement standard deviations) served as a quality score for the used net flux sample. As the isotopomer model contains non-linear equations, there is no guarantee of reaching a global optimum within reasonable time and efforts (Antoniewicz et al, 2006). Thus, we first performed optimisations with different combinations of maximum number of solver iterations and number of random starting points. Using the [U-^13^C] data set as a test case, we found that 200 iterations and 50 starting points yielded the lowest optimisation residuals in acceptable amounts of time (Fig. S4a,b). Using these numbers of iterations and starting points, in independent optimisations, we fitted the model to the ^13^C labelling data, with 1000 different net flux samples from the feasible TSM flux space as input (Supplementary Data File 4).

Prior to using the net flux samples determined with the sampling approach, we had also attempted to fit the isotopomer model to labelling data without providing net flux solutions, by just using the upper and lower net flux bounds obtained from the variability analysis of the TSM model (cf. Fig. S1a). Here, optimisations stopped because of very slow progression in the objective function even before reaching the maximal number of iterations, with residual values in the order of 10 000. The limited progression in the optimisation suggests that the solution space, as constrained by the net flux bounds, is too large for the solver to find any solution with a good fit. Of note, even if the fittings were successful, the net fluxes obtained in such a manner would not necessarily comply with thermodynamics, as there is still the possibility for thermodynamically infeasible loops. These findings underline the need for the approach outlined above, which makes use of thermodynamically feasible net flux samples.

Next, we evaluated whether net flux samples from the stoichiometric model (SM) fitted to physiological data, would be sufficient to provide a good fit of the isotopomer model to the ^13^C data. For sampling, we used the optGpSampler (Megchelenbrink et al, 2014) implemented in the COBRA Toolbox (Heirendt et al, 2019). Of note, also these net flux samples do not necessarily comply with the laws of thermodynamics. Using these net flux samples in the fitting of the isotopomer model to the [U-^13^C] data set, we found that the residuals were between 40000 to 60000 (Fig. 5a), with several isotopomer abundances not fitting well (Fig. 5b). In contrast, fitting the isotopomer model to the same data while providing the thermodynamically-constrained net flux samples as input, we obtained residuals in the order of 2000 (Fig. 5a), and the isotopomer abundances were much better fitted (Fig. 5b). The higher residuals obtained with the SM net flux samples, compared to the TSM net flux samples, indicate that the SM flux solution space is too large, resulting in a small probability for the optimiser to find good starting points. Thus, thermodynamics together with metabolome and physiological data reduce the original SM flux solution space to a much smaller sub-space where the actual fluxes lie, facilitating the fitting of the ^13^C labelling data.

**Figure 5:**
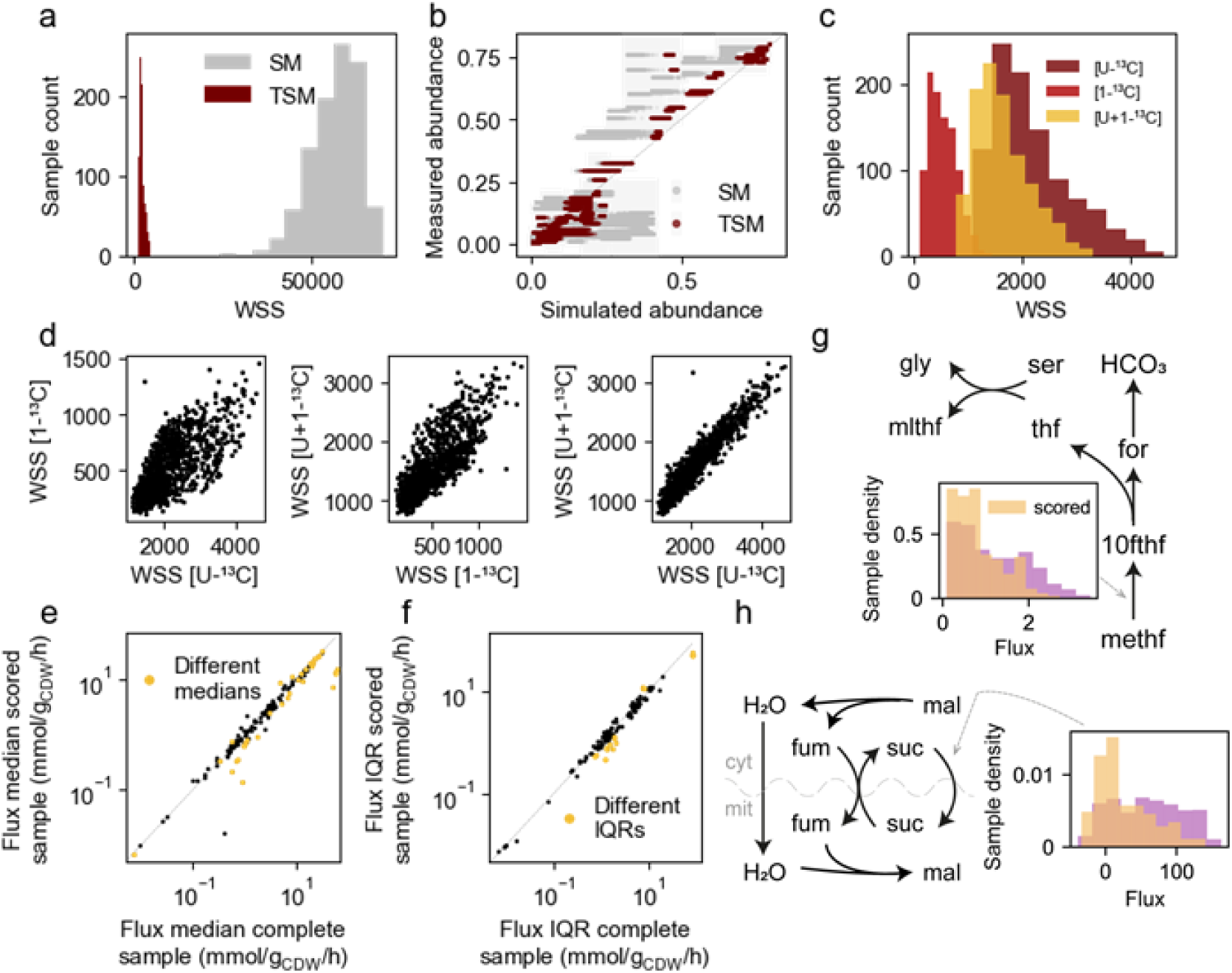
Scoring net flux samples with ^13^C labelling data. (a) Optimisation residuals of the fitting of the isotopomer model to ^13^C labelling data with net flux samples from the stoichiometric (SM) and thermodynamic model (TSM), where WSS stands for a weighted sum of squares, referred to in the text as residual. (b) Measured versus simulated isotopomer abundances of all measured amino acids, obtained by fitting the SM and TSM models with labelling data with the respective net flux samples fixed, where the background boxes (light grey) represent the standard deviation of the measurements and the diagonal dashed line represents the same value for x and y-axis. (c) Optimisation residuals obtained by fitting the isotopomer model to [U-^13^C], [1-^13^C] and both data sets simultaneously [U+1-^13^C] with TSM net flux samples. [U-^13^C] represents data from a labelling experiment with a mix of 80% unlabelled glucose and 20% fully labelled glucose and [1-^13^C] represents data from a labelling experiment with 100% glucose labelled in the first carbon atom. (d) Scatterplots of residuals (WSS) for the three different runs of fittings with [U-^13^C], [1-^13^C] and [U+1-^13^C]. In (c) and (d), the value of WSS of fitting with [U+1-^13^C] data was divided by two because it contains double the number of measurements (measurements of [U-^13^C] and [1-^13^C]) compared to fitting only [U-^13^C] or [1-^13^C]. (e) Flux medians of a set of TSM flux samples (i.e. 1000 net flux samples obtained from Parallel Tempering) versus the subset consisting of the top 20% samples from the scoring with the [U+1-^13^C] data (i.e. 200 net flux samples). To determine the medians with statistically significant differences between the set and the subset of scored samples, a statistical analysis was performed. Several random subsets of size 200 were drawn from the set of net flux samples and a “null” distribution of medians was built from the random subsets. If the median of the flux distribution (from the scored set of samples) of a particular reaction was outside the interval defined by the 0.01% and 99.99% percentiles of the “null” distribution, the flux distribution of that reaction was considered statistically different (yellow in the plot) compared to the set of samples before scoring. (f) Flux interquartile ranges (IQRs) of a set of TSM flux samples (i.e. 1000 net flux samples obtained from Parallel Tempering) versus the subset consisting of the top 20% samples from the scoring with the [U+1-^13^C] data (i.e. 200 net flux samples). The same statistical analysis as for the medians was applied to the IQRs and the statistically different IQRs are represented in yellow in the plot. (g), (h) Sub-networks of folate metabolism and malate exchange respectively. Reactions in these sub-networks were affected by the scoring of the sample, as in they were flagged with different medians in the statistical analysis performed in (e). One example histogram is shown for each subnetwork with a flux distribution for the complete sample (pink) and the scored sample (yellow), while the rest of the fluxes are not shown as they have similar shapes to the ones presented. The direction of arrows is the direction of the median of the flux. All data and analyses mentioned here refer to the WT. cyt: cytosol, mit: mitochondria, gly: glycine, ser: serine, mlthf: 5,10-methylenetetrahydrofolate, thf: tetrahydrofolate, HCO3: bicarbonate, for: formate, 10fthf: 10-formyltetrahydrofolate, methf: 5,10-methenyltetrahydrofolate, mal: malate, fum: fumarate, suc: succinate.

Ultimately, our goal was to fit both ^13^C-labelling data sets [U+1-^13^C] simultaneously, as such integrated analyses were found to yield improved results (Antoniewicz, 2015). However, before we did so, we asked whether the two data sets provide diverging information on the scoring of net flux samples. Comparing the optimisation residuals obtained by fitting the model to [U-^13^C] and [1-^13^C] individually and simultaneously, using wildtype net flux samples we found that the [1-^13^C] data yields lower residuals compared to the [U-^13^C] data (Fig. 5c). When fitting the model to both labelling data sets simultaneously ([U+1-^13^C]), we obtained intermediate residuals (Fig. 5c). Plotting residuals of individual net flux samples obtained from fittings to [U-^13^C], [1-^13^C] and [U+1-^13^C] against each other, we found that, globally, the residuals correlate. This indicates that for each net flux sample the different labelling data sets convey similar information (Fig. 5d, Fig. S4c).

To determine the constraining effect of the [U+1-^13^C] labelling data on the TSM net flux samples, we selected the 20% best fitting net flux samples, i.e. those with lowest residuals, from a set of TSM net flux samples (of size 1000). We first generated different random subsets of the net flux samples (of size 200). From these random subsets, we calculated the empirical “null” distributions of flux medians and interquartile ranges (IQR) for each reaction. Next, we identified the reactions for which the median and IQR of the top 20% samples from the scoring with the [U+1-^13^C] data was outside the interval defined by the 0.01% and 99.99% percentiles of the “null” distribution. We considered that for these reactions there was a significant change in median or IQR compared to the distribution of outcomes that would be obtained by chance alone. We found that the ^13^C labelling data had an effect on the medians of 40 reaction fluxes (Fig. 5e) and the IQRs of 26 reaction fluxes (Fig. 5f). Nine of these reactions are connected in two subnetworks with more than two reactions, indicating that addition of the 13C data constrains not isolated reactions but rather sub-networks. One of these subnetworks is related to folate metabolism (Fig. 5g) and the other to malate exchange from the cytosol to the mitochondria (Fig. 5h).

Thus, at this point we can conclude the following: with metabolic networks of realistic size and with full reaction reversibility allowed, (i) net flux samples from a stoichiometric model constrained by physiological data only deliver suboptimal fits, (ii) fitting an isotopomer model to ^13^C labelling data and net flux bounds (obtained from TSM) with net fluxes as free variables is also not able to reach a good fit as the problem is too unconstrained. Both methods would also not be guaranteed to provide thermodynamically feasible net flux samples. In contrast, the net flux samples from the TSM yielded improved residuals in the isotopomer model fits compared to (i) and (ii). Overall, while the addition of ^13^C labelling data constrained fluxes in certain sub-networks, our analysis showed that the ^13^C addition only introduces small changes in the flux samples constrained by thermodynamics, stoichiometry, metabolome, and physiological data.

### Further constraining with the flux force relationship

Up to this point, we have constrained the flux solution space with thermodynamics, stoichiometry, metabolome and physiological data. However, there is the possibility to add another constraint that we have not exploited yet. Specifically, given a TSM net flux sample, we can also obtain Δ_*r*_*G* samples for all reactions by performing an additional sampling of Polytope B. This piece of information offered the opportunity of adding an additional constraint to the fitting of the isotopomer model via the so-called flux force relationship (Rastogi, 2007). This relationship links the Δ_*r*_*G* of a reaction with the ratio between its forward (*v*^+^) and backward fluxes (*v*^−^), according to

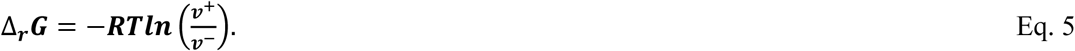

However, the flux force relationship, as defined in Eq. 5, only holds for simple reaction kinetics (Beard & Qian, 2007;Wiechert, 2007). For more complex reaction mechanisms, the equality turns into an inequality (Beard & Qian, 2007), where

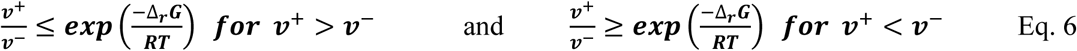

These inequalities can be rewritten in terms of the net and labelling exchange fluxes, which are the variables in the isotopomer model we fit to the ^13^C labelling data, yielding

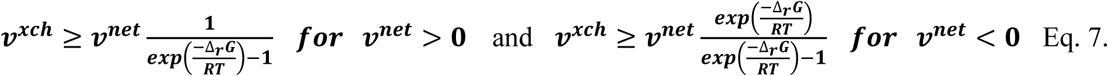

These relationships, together with sampled values of Δ_*r*_*G* for each reaction, can provide us with lower bounds on the labelling exchange fluxes, ***v*^*xch*^**, which we could exploit when fitting the ^13^C labelling data. To obtain Δ_*r*_*G* samples, we sampled the concentration polytope (i.e. polytope B in Fig. 3a) for each net flux sample with the HR algorithm. For each net flux sample, this approach generated an empirical distribution of Δ_*r*_*G* values for each reaction. From these distributions, we selected the Δ_*r*_*G* values that resulted in the least conservative (i.e. lowest) bound on the labelling exchange flux. Using these Δ_*r*_*G* values and the flux force inequality (Eq. 7), we calculated the bounds for the labelling exchange fluxes, which we then used in the fitting of the isotopomer model to the ^13^C data [U+1-^13^C], with the respective net flux samples as input. We did not define labelling exchange flux bounds for transport reactions with more than one variant, because, as mentioned above, different transporter variants were not included in the isotopomer model.

Consistent with the presence of additional constraints, the optimisation over the labelling exchange fluxes produced residuals that spread over a broader range of values and were on average higher (Fig. 6a) than without the constraints. Remarkably, the best fitting net flux samples obtained with the flux force constraint had residuals that were almost as low as the residuals without (Fig. 6a). Plotting the residuals corresponding to net flux samples obtained with and without flux force constraints against each other, we found that the best-fitting net flux samples have residuals in the order of 1000 in both cases. Moreover, there is a good correlation between residuals corresponding to net flux samples scored with and without the flux-force constraints (Fig. 6b). These findings suggest that, while the addition of the flux force relationship yields a comparable relative ranking of the net flux samples, at the same time, it allows for better discrimination between flux samples.

**Figure 6:**
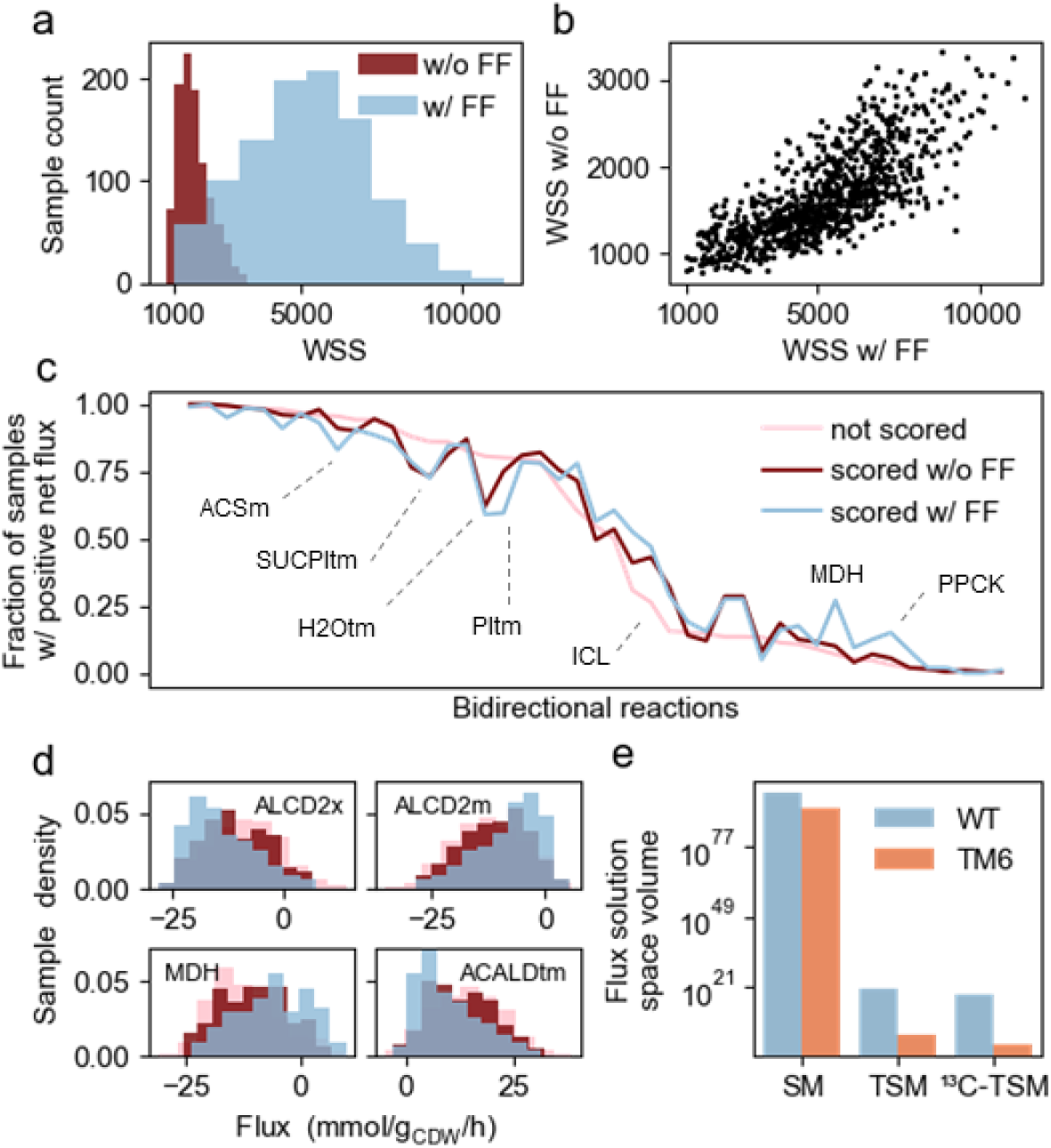
Further constraining the flux solution space with the flux force relationship and Gibbs energies of reaction. (a) Optimisation residuals obtained by fitting the isotopomer model to [U+1-^13^C] data with (w/ FF) and without (w/o FF) flux force constraint. The flux force constraint is applied to the fitting problem as a lower bound on the labelling exchange fluxes through Eq. 7 using limits from sampled Gibbs energies of reaction (Δ_*r*_*G*). WSS stands for a weighted sum of squares, referred to in the text as residual. (b) Scatter plot of residuals (WSS) of the fit with (w/ FF) and without (w/o FF) flux force relationship. In (a) and (b), WSS values are represented as half of their original value, so that the values are consistent with those in Fig. 5c and d. (c) The fraction of samples where bidirectional reactions have a positive net flux in the three differently scored samples. Samples without scoring, pink; scored samples without FF, dark red; scored samples with FF, blue. Bidirectional reactions are reactions for which we have net flux samples in both directions. The reactions are here sorted along the x-axis according to the fraction of the non-scored sample (pink), shown on the y-axis. Some of the reactions with the highest difference among differently scored samples are identified with their names. ACSm: mitochondrial acetyl-CoA synthetase, SUCPItm: combined transport of succinate and phosphate across the mitochondrial membrane, H2Otm: water transport across the mitochondrial membrane, PItm: one of the variants of phosphate transport, ICL: isocitrate lyase, MDH: cytosolic malate dehydrogenase, PPCK: phosphoenolpyruvate carboxykinase. (d) Flux distribution of the four reactions identified as having statistically different flux distributions between the 20% best scoring net flux samples from the fit with (w/ FF) and without (w/o FF) flux force constraint, as assessed by a Kolmogorov-Smirnov test applied to the flux distributions of each reaction, with a significance level of 0.001. Colours as in (c). ALCD2x: cytosolic ethanol dehydrogenase, ALCD2m: mitochondrial ethanol dehydrogenase, MDH: cytosolic malate dehydrogenase, ACALDtm: acetaldehyde transport across the mitochondrial membrane. (e) Flux solution space volume for WT (blue) and TM6 (orange) calculated with Expectation Propagation for models with increasing levels of constraints. This volume is a relative measure used to compare different solution spaces but with no physical meaning on its own. SM: stoichiometric model and physiological data, TSM: thermodynamic/stoichiometric model with metabolome and physiological data, ^13^C-TSM: 20% best samples from TSM according to fitting to [U+1-^13^C] data using flux force constraint.

Next, we asked which reaction fluxes would assume different values in the two scorings, i.e. with and without flux force relationship constraint. For a first assessment of the effect of the addition of the flux force relationship on reaction directions, we plotted the fraction of net flux samples where the reaction goes into the positive direction (according to the specification in the stoichiometric model) over the total number of independent reactions (i.e. reactions that are not in the same linear pathway). We did this for the complete set of net flux samples before scoring, for the 20% best-fitting net fluxes after the ^13^C fit, and the 20% best-fitting net fluxes after the ^13^C fit with flux force relationship constraint. Here, highlighting the effect of the flux force relationship constraint, we found that for several reactions this fraction changes (Fig. 6c). To compare fluxes between the two samples (i.e. the 20% best scoring net flux samples with and without flux force constraint), we used a Kolmogorov-Smirnov test. Here, focussing on the 20% best scoring net fluxes from the fits with and without flux force constraint, we found four reactions with statistically different flux distributions (Fig. 6d). Thus, the addition of the flux force as an inequality constraint to the model fitting with ^13^C data allowed for better but relatively similar discrimination of the net flux samples, while having only a small effect on the individual flux distributions.

To assess the size of the flux solution space after each of the above-applied steps, we approximated the distributions of the generated flux samples by multivariate Gaussian distributions, from which we then estimated the volume of the flux solution space under different sets of additional constraints, using Expectation Propagation (Braunstein et al, 2017). We found that the volume of the flux solution space constrained by the stoichiometry and physiological data is 10^98^, the one additionally constrained by thermodynamics and metabolome data is 10^20^, and the one corresponding to the 20% best-fitting samples after the fit to the [U+1-^13^C] data with flux force constraint is 10^18^ (Fig. 7e). Similar reductions also occur with TM6 (Fig. 7e). Thus, each step reduces the size of the flux solution space, with the biggest drop occurring in the step where thermodynamics and metabolome data are added.

**Figure 7:**
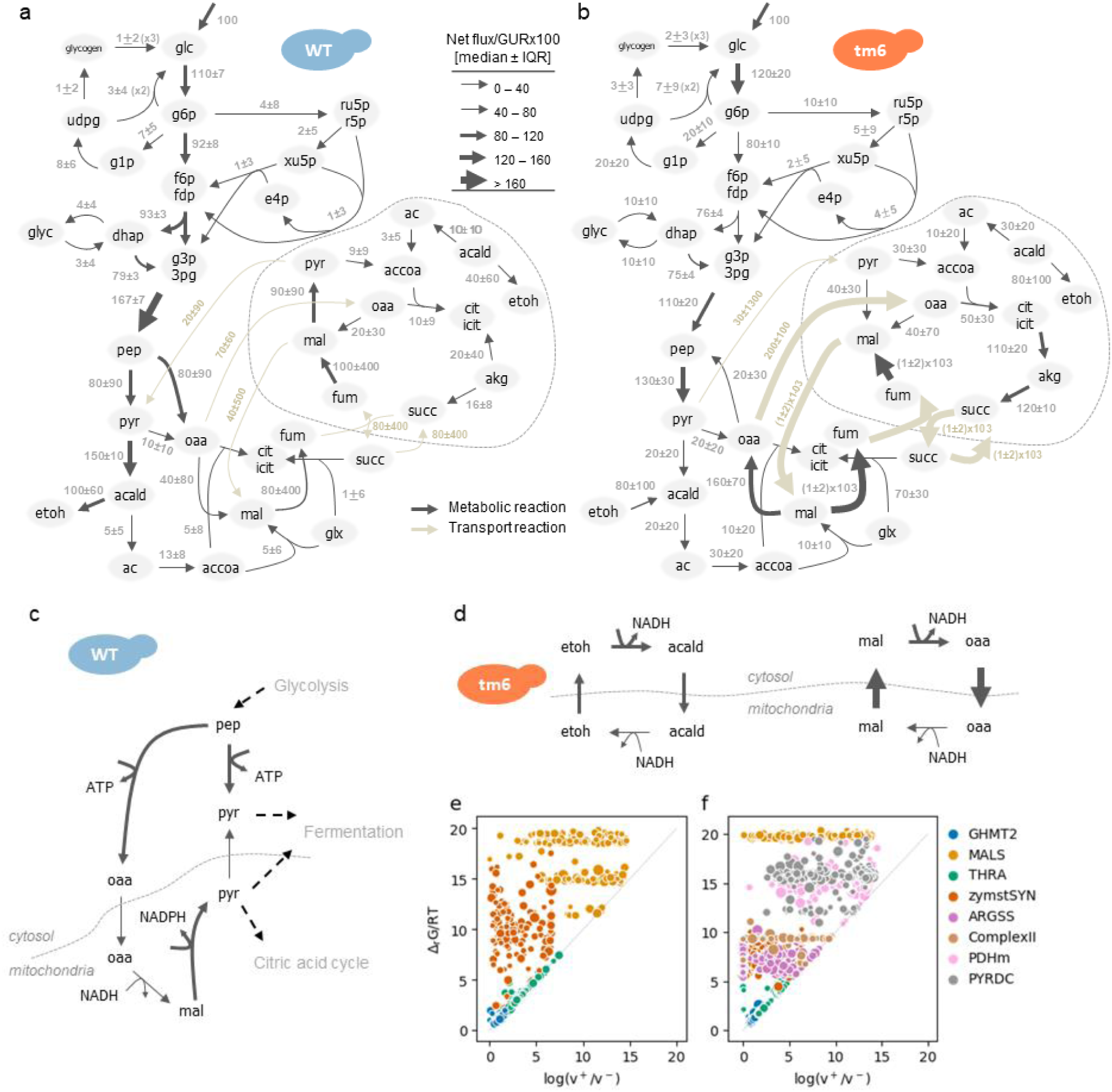
Metabolic fluxes of yeast under glucose conditions. (a),(b) Visualisation of net flux distributions in the central metabolism of WT (a) and TM6 (b). Grey numbers represent median and interquartile range of net flux distributions where 20% best fitting samples were chosen based on residuals of ^13^C fitting with flux force constraint, values scaled to 100 of glucose uptake (GUR). Arrow thickness and direction represent the median of net flux distributions on the same scale as grey numbers. Arrows of different colour represent transport reactions between cytosol and mitochondria. The reactions within the dashed line occur in the mitochondria. (c) Zoom in of metabolic network visualising the flux around the anaplerotic/gluconeogenic reactions in WT. (d) Two NADH shuttles running for TM6 but not WT. In (c) and (d) arrows represent reactions, arrow direction represents the direction of median flux and arrow thickness represents flux median value with same scale as in (a) and (b). (e,f) Relationship between the ratio of Gibbs energy of reaction over gas constant times temperature (Δ_*r*_*G*/RT), and the logarithm of the ratio between forward and backward flux (log(v^+^/v^-^)) for the determined labelling exchange fluxes in WT (e) and TM6 (f). Colours denote different reactions, for which there are several values corresponding to several net flux samples of the 20% best samples according to the residuals of ^13^C fitting with flux force constraint. Point size is inversely scaled with the fitting residual: a big point has a better fit to the data than a small point. The Δ_*r*_*G* values used in these plots are the same used to impose the lower bounds on the labelling exchange fluxes. Hence, all values are forced above the diagonal, which represents the flux-force equality (Eq. 5). Transport reactions that have different stoichiometric variants in the model are not represented here as we could not relate one value of flux to the several values of sampled Δ_*r*_*G* for the different variants. This is the case for the mitochondrial glyoxylate transport, which was the only transporter for which we could determine the labelling exchange flux, both in WT and TM6. GHMT2: glycine hydroxymethyltransferase (EC 2.1.2.1), MALS: malate synthase (EC 4.1.3.2), THRA: threonine aldolase (EC 4.1.2.5), zymstSYN: zymosterol synthesis, ARGSS: argininosuccinate synthase (EC 6.3.4.5), Complex II: succinate dehydrogenase (EC 1.3.5.1), PDHm: pyruvate dehydrogenase (EC 1.2.4.1), PYRDC: pyruvate decarboxylase (EC 4.1.1.1).

### Flux distributions unravelled

Applying the different sets of data and constraints, we obtained distributions for each reaction flux of budding yeast under two important metabolic conditions (aerobic fermentation & and respiration, Supplementary Data File 8) that are consistent with stoichiometry, thermodynamics, physiological, metabolome and ^13^C labelling data. To our knowledge, these are the most unbiased flux distributions that were determined so far, i.e. with a large metabolic network model (258 reactions) and without the use of any *a priori* constraints on reactions’ directions and reversibility. To gain insight from these distributions, we visualised them on a metabolic map with medians and IQRs, for both the WT and TM6 (Fig. 7a,b).

Here, we found that the ratio between the glycolytic and pentose phosphate pathway flux in the WT is 0.03 ± 0.08 (median ± IQR), and 0.11 ± 0.14 in the TM6. These values are in good agreement with values from the literature, i.e. 0.03 and 0.22, respectively (Gombert et al, 2001). Yet, consistent with the fact that current ^13^C flux analyses likely use over-constrained models, we found flux patterns that are different from the currently published ones. Specifically, we found that in the WT there is an almost cyclic flux from phosphoenolpyruvate via oxaloacetate (catalysed by phosphoenolpyruvate carboxylase, PCK1) to mitochondrial oxaloacetate, mitochondrial malate, mitochondrial pyruvate and back to cytosolic pyruvate (Fig. 7c), overall converting mitochondrial NADH into NADPH as a source of mitochondrial NADPH for biosynthesis. This flux accounts for about 50% of the phosphoenolpyruvate-to-pyruvate conversion, the remainder of which is catalysed by pyruvate kinase (Fig. 7a). In TM6, the malic enzyme and the phosphoenolpyruvate carboxylase go in the opposite directions (Fig. 7b), following the common notion on these reactions’ directions (Zelle et al, 2010). While genetic data in principle argues against a flux from phosphoenolpyruvate to oxaloacetate (Brewster et al, 1994; Stucka et al, 1991), it was shown that the PCK1 reaction, at reduced pyruvate kinase activity, can support an anaplerotic function of PCK1 (Zelle et al, 2010), which is the activity that we find here with our unbiased flux determination approach. The directional operation of these fluxes as we find them in the WT has so far remained unknown, likely because current ^13^C flux analyses employ metabolic networks that are over-constrained with regards to reaction directions.

In the absence of a cytosolic-mitochondrial NADH transporter, several NADH shuttles are known to be present in *S. cerevisiae* to ‘shuttle’ NADH in and out of the mitochondria (Bakker et al, 2001). According to our estimated flux distributions, two NADH shuttles are operative in TM6, i.e. the ethanol-acetaldehyde shuttle and the malate-oxaloacetate shuttle (Fig. 7d). Both shuttles operate in the direction of shuttling NADH from the cytosol into the mitochondria. In the WT, based on the median flux values, both shuttles are not operative although we cannot exclude that the malate-oxaloacetate shuttle is active, given that the flux distribution of the cytosolic malate dehydrogenase reaction spans from positive to negative values in the WT (Fig. S5).

Finally, we asked whether we could pinpoint magnitudes of labelling exchange fluxes. To test this and to identify labelling exchange fluxes, we focussed on the labelling exchange fluxes from the 20% best scoring fits when we used the flux force constraint. For each labelling exchange flux (of each optimisation, Supplementary Data File 9) we determined the respective standard deviation from the Jacobian of the measurements with respect to the free variables (i.e. all labelling exchange fluxes) and the measurement covariance matrix. For the labelling exchange flux of each reaction, we then determined the fraction of optimisations in which the standard deviation was below an empirical value here defined as 200 mmol/gDW/h (Fig. S6a,b). We considered labelling exchange fluxes as ‘determined’ when they had at least 50% of the solutions within this standard deviation threshold. Based on this criterion, we identified the labelling exchange fluxes of five reactions for WT and of ten fluxes for TM6.

After obtaining, for several reactions, net fluxes, flux ratios and labelling exchange fluxes, as well as Δ_*r*_*G* (Fig. S6c,d), we finally asked for which of these reactions the flux-force relationship would be an equality (Eq. 5). To this end, we plotted the highest absolute value of the Δ_*r*_*G* distribution (i.e. the one used for the flux force constraint) against the logarithm of the forward and backward flux ratio for each net flux sample of WT (Fig. 7e) and TM6 (Fig. 7f). Here, we found that, for both strains, only glycine hydroxymethyltransferase (GHMT2) and threonine aldolase (THRA) strictly follow the flux-force equality, suggesting that these reactions follow a simple reaction mechanism, where the flux force equality strictly holds.

## DISCUSSION

With this work, we improved our ability to determine intracellular metabolic fluxes. We accomplished this by developing a novel pipeline, in which we use physiological, metabolome and ^13^C labelling data together with a thermodynamic and stoichiometric metabolic network model, in an approach that goes a step beyond earlier work (Park et al, 2019). Our pipeline generates intracellular flux estimates for a large network model without any assumptions on reaction direction or labelling exchange fluxes. Furthermore, providing an alternative to a Bayesian approach to estimate flux uncertainties (Theorell et al, 2017), with our approach, we obtain not just one flux solution, but also statistical estimates of the uncertainty of each reaction flux. Because a simultaneous computational integration of these three different data types (i.e. physiological, metabolome, and ^13^C data) is likely not possible due to the complexity introduced by the thermodynamics and the isotopomer model, we accomplished the integration through a three-step approach. In the flux distributions, which we generated for budding yeast grown under fermentative and respiratory metabolic modes, we discovered novel flux patterns that until now had remained unknown, likely due to the assumptions made in the current ^13^C metabolic flux analysis investigations.

Our method required an approach to comprehensively sample the complex non-convex solution space of flux, metabolite concentration, and Δ_*r*_*G* constrained by the thermodynamic and stoichiometric model and the metabolome and physiological data. As none of the existing samplers (Binns et al, 2015;Chaudhary et al, 2016;Saa & Nielsen, 2016) was able to accomplish this, we developed a new approach to efficiently sample this space. Our sampling approach comprises several steps, including the splitting of the solution space into convex polytopes, the scaling of the flux polytope and its division in sectors, and the application of Hit-and-Run in a Parallel Tempering framework. We envision that our sampling approach, standing by itself, will be useful for the community to sample complex metabolic flux solution spaces.

Compared to current ^13^C metabolic flux analyses, our approach provides less biased flux estimates. First, often only small metabolic networks are used or the employed networks do not describe the complexity of compartmentalised metabolism in detail. Contrarily, we apply our method to a metabolic network of 258 reactions, with 148 independent fluxes including central carbon metabolism and different variants of transmembrane transport between the cytosolic and mitochondrial compartment. Second, reactions are often assumed to have particular directions, which allowed for ^13^C metabolic flux analysis at genome-scale (Basler et al, 2018;Gopalakrishnan & Maranas, 2015). We do not limit any reaction direction *a priori* and instead let the data and the model decide, based on thermodynamics. With this approach, we found several reactions with opposite directions in the two metabolic modes and we found previously unknown flux patterns. Third, assumptions are also made with regards to the occurrence and/or magnitude of labelling exchange fluxes. For instance, it is often assumed that there is no labelling exchange flux for particular reactions or the flux-force equality is applied to constrain the labelling exchange fluxes (Beard & Qian, 2005;Noor et al, 2014). Our work shows that the use of the flux force relationship as an equality may only apply for some reactions (Figs. 7e,f), in line with the finding that for complex enzymatic reactions flux force relationship should only be used as inequality (Beard & Qian, 2007). With our method, we provided an approach, with which none of these assumptions needs to be made to estimate metabolic fluxes.

Further, our work shows that once the flux space is constrained by the combined thermodynamic-stoichiometric metabolic model and metabolome and physiological data, then the addition of ^13^C labelling data from amino acids has only a small additional constraining effect on the overall flux solution space. For example, we found that the ^13^C labelling data only decreased the flux variability of a handful of reactions (Fig. 5f). Different fractions of best fitting net flux samples (i.e. best 5%, 10%, etc.) also yielded very similar distributions of reaction flux medians and IQRs (Fig. S7a,b). In contrast, when we fitted the ^13^C labelling data to net flux samples generated from only a stoichiometric model fitted to physiological data, the ^13^C optimisations resulted in very high residuals (Fig. 5a), which indicates that more diverse net flux samples (i.e. not constrained by thermodynamics and metabolome data) can indeed yield much worse fittings. Overall, these findings suggest that metabolome and physiological data together with a combined thermodynamic-stoichiometric model constrain the flux solution space already so much, that ^13^C labelling data determined in amino acids can hardly contribute any further. Yet, the situation will likely change when ^13^C labelling data in metabolites of central carbon metabolism, as acquired with novel experimental techniques (Feith et al, 2019;Jaiswal et al, 2018;Mairinger et al, 2015), or non-carbon isotope tracers (Xu et al, 2020) are exploited as well.

With our approach, we can now generate high-quality, unbiased flux estimates. This will allow the identification of flux differences between diverse metabolic modes and support metabolic engineering efforts. Further, the fluxes that we could infer in an unbiased manner, and in particular the new flux patterns we uncovered, provide new insights on the intracellular yeast physiology, which requires that some of the current notions on metabolic flux distributions in yeast need to be revisited. Finally, while according to our findings physiological and metabolome data are sufficient to obtain excellent flux estimates for conditions where yeast is grown on minimal media, we envision that ^13^C data, as applied in this work, can additionally contribute to unravel fluxes also under conditions where cells grow on complex media (Schwechheimer et al, 2018).

## METHODS

### Yeast strains and maintenance conditions

The haploid, prototrophic *Saccharomyces cerevisiae* strain KOY PK2-1C83 (relevant genotype: wild type) and its derivative KOY TM6*P (relevant genotype: chimeric *HXT1*-*HXT7* glucose transporter as sole hexose transporter) (Elbing et al, 2004) were used in this study. The medium used for yeast strain maintenance was YPD, which contained 2% [v/w] peptone, 1% [v/w] yeast extract and 2% [v/w] glucose.

### Shake flask cultivation and labelling experiments

All cultivations were performed at 30°C and cultures were shaken at 300 rpm. The minimal medium was composed as follows (Verduyn et al, 1990): 5 gL^-1^ (NH_4_)_2_SO_4_, 3 gL^-1^ KH_2_PO_4_, 0.5 gL^-1^ MgSO_4_·7H_2_O, 1.5 mgL^-1^ EDTA, 4.5 mgL^-1^ ZnSO_4_·7H_2_O, 0.3 mgL^-1^ CoCl_2_·6H_2_O, 1 mgL^-1^ MnCl_2_·4H_2_O, 0.3 mgL^-1^ CuSO4·5H_2_O, 4.5 mgL^-1^ CaCl_2_·2H_2_O, 3 mgL^-1^ FeSO_4_·7H_2_O, 0.4 mgL^-1^ Na_2_MoO_4_·2H_2_O, 1 mgL^-1^ H_3_BO_3_, 0.1 mgL^-1^ KI, 0.05 mgL^-1^ D-Biotin, 1 mgL^-1^ Ca-pantothenate, 1 mgL^-1^ Nicotinic acid, 25 mgL^-1^ m-Inositol, 1 mgL^-1^ Pyridoxine HCl, 0.2 mgL^-1^ p-Aminobenzoic acid, 1 mgL^-1^ Thiamine HCl, and 10 gL^-1^ D-glucose. The medium was buffered at pH 5 with 10 mM KH-phthalate.

To ensure exponential growth of the cells, and thus a metabolic steady state, from which the samples were taken, two pre-culturing steps were carried out before the main culture. The first pre-culture of 3 mL was grown to stationary phase and used to inoculate the second preculture at an OD_600_ of 0.1 - 0.2. The cultivation of the second inoculum was carried out in 500 mL Erlenmeyer shake-flask containing 50 mL of minimal medium. From the mid-exponential growth phase of this second culture with an approximate cell density ranging between 10^7^ and 10^8^, the main culture was inoculated. To avoid excessive carryover of unlabelled cells from the pre-culture, the inoculum of exponentially growing cells in the main culture was kept below 10^5^ cells mL^-1^, which were transferred from the second culture. The second pre-cultures and main cultures were grown using identical cultivation conditions, i.e. minimal medium supplemented with glucose, a temperature of 30 °C and continuous shaking at 300 rpm. After the inoculation of the main culture, the WT and TM6 strains were cultivated for 8h and 12h overnight. In this unobserved period, the main cultures of the WT and TM6 reached in the morning an approximate cell count of 10^5^ cells mL^-1^ and 10^6^ cells mL^-1^, respectively. After this unobserved period, the cell counts were then measured every 1.5 hours until the mid-exponential growth phase of the strains, when the WT and TM6 reached again an approximate cell density ranging between 10^7^ and 10^8^ cell mL^-1^.

For quantification of the metabolite uptake and consumption rates, the two strains were cultivated in three independent cultivations in 500 mL Erlenmeyer flask containing 50 mL of minimal medium supplemented with 10 gL^-1^ glucose. The ^13^C labelling experiments were done in duplicates in a 100 mL Erlenmeyer flask with 10 mL minimal medium and 10 gL^-1^ of a defined mixture of glucose isotopomers. The two glucose isotopomer mixtures used were composed of either 100% [1-^13^C] glucose with the ^13^C label at the first carbon atom, or 80% unlabelled and 20% uniformly [U-^13^C] labelled glucose, as described earlier (Zamboni et al, 2009). The labelled glucose was only present in the last main culture. As we inoculated this last culture only with around 50’000 cell mL^-1^ and harvested cells at 10^7^ and 10^8^ cell mL^-1^, the effect of unlabelled biomass is marginal.

### Quantification of biomass dynamics

Cell counts were measured in a Accuri C6 flow cytometer. Before measuring, the culture was diluted to a cell count concentration below 10^6^ cells mL^-1^ with minimal medium, and the cell counts were measured in a total volume of 20 µL, with the fluidics setting set to ‘medium’. The FSC-H thresholds was set to 80 000 to cut off most of the electronic noise. The Accuri CFlow Plus software was used for data analysis. The cell count was determined approximately every 1.5 hours. At the end of the culture, the yeast dry mass was determined by filtering a volume of 40 mL and 35 mL culture broth of the WT and TM6 through pre-weighed nitrocellulose filters with a pore size of 0.2 µm. Filters were washed once with distilled water and kept at 80°C for two days. Afterward, they were weighted again. The cell dry mass at every measurement point was re-calculated from the cell count and the dry mass cell count/ratio using the dry mass cell count/ratio obtained at the end of the culture.

### Quantification of extracellular metabolite dynamics

Toward determining the glucose uptake and pyruvate, glycerol, acetate, and ethanol production rates, samples of supernatant were taken from three independent cultivations in 50 mL. The concentrations of glucose, pyruvate, glycerol, acetate, and ethanol in the supernatant were determined by HPLC (Agilent, 1290 LC HPLC system) using a Hi-Plex H column and 5 mM H_2_SO_4_ as eluent at a constant flow rate of 0.6 mL min^-1^. The column temperature was kept constant at 60°C and a volume of 10 μL was injected for analysis. Substrate concentrations were detected with refractive index and UV (constant wavelength of 210 nm) detection. The chromatogram integration was done with Agilent Open Lab CDS software. Metabolite concentrations were determined using external calibration with a serially diluted standard, which included all metabolites, relevant for the various conditions. The standards covered the metabolites’ concentration range, from the start until the end of the batch cultivation in glucose. The generation of the standard curve was done before the sample analysis and thereafter updated with control standards, which were included in every sample run.

### Determination of physiological rates

Exchange fluxes for glucose, ethanol, acetate, glycerol, pyruvate, and biomass were estimated by fitting the concentration time courses to an ordinary differential equation model (ODE) assuming exponential growth and constant yields in the batch culture. The ethanol evaporation was included with a first-order rate in the ODE model with the evaporation rate proportional to the ethanol concentration. The evaporation rate was pre-determined from an independent experiment, where the ethanol concentration was followed in 50 mL minimal medium supplemented with an initial ethanol concentration of 4 gL^-1^ in 500 mL Erlenmeyer flasks shaken under cultivation conditions, i.e. 300 rpm and 30°C. The evaporation constant was estimated to be 0.021 h^-1^ by fitting an exponential function to the measured ethanol time course. We used a previously described model (Litsios et al, 2019) and gPROMS Model Builder v.4.0 (Process Systems Enterprise Ltd.) for parameter estimation.

### Quantification of mass isotopomer patterns in ^13^C labelling experiments

Three different types of labelling experiments were performed in duplicates, each with a total glucose concentration of 10 g L^-1^. The first set of experiments contained only unlabelled glucose to determine the naturally occurring isotopomer ratios. The second set of cultures contained [1-^13^C] glucose (Omicron, GLC-018). The third set of cultures contained a mixture of 80% unlabelled and 20% uniformly labelled [U-^13^C] glucose (Buchem, CLM-1396). From each flask, three 1 mL samples of culture broth were harvested at an approximate cell count of 10^7^ cell mL^-1^, corresponding to the early mid exponential phase. After harvest, samples were centrifuged and the supernatant was discarded. The cell pellets were washed once in 1 mL 0.9% (w/v) NaCl and frozen at -80°C until further use. Further sample preparation for isotopomer analysis of proteinogenic amino acids was performed as previously done (Zamboni et al, 2009). Specifically, the pellets were hydrolysed for 24 h in 200 µL 6M HCl to release the proteinogenic amino acids. The hydrolysates were dried at 105°C and the dried pellets resuspended in 20 µL of dimethylformamide (DMF, Sigma Aldrich). The amino acids were derivatised adding 20 µL of N-tertbutyldimethylsilyl-N-methyltrifluoroacetamide (TBDMSTFA; Sigma Aldrich) for 1h at 85°C.

The derivatised sample preparations were analysed in triplicate by GC-MS analysis according to the method described earlier (Zamboni et al, 2009) 1 µL of derivatised sample was injected into the Agilent 7890A Series GC with inert MSD/DS Std Turbo EI System. The TBDMS derivatives were separated on the DB-5MS GC Column of Agilent (Product number 122-5532) applying conditions as previously published (Zamboni et al, 2009). The analysed proteinogenic amino acids were alanine, glycine, valine, leucine, isoleucine, proline, methionine, serine, threonine, phenylalanine, aspartate, glutamate, lysine, histidine, and tyrosine. Mass spectrometric raw data of the hydrolysed proteinogenic amino acids were analysed using MZmine2 (Pluskal et al, 2010). The assignment of peaks in the total ion chromatogram to specific proteinogenic amino acids was done based on retention times taken from previous published data (Zamboni et al, 2009). Towards determining the mass isotopomer patterns of an amino acid, we defined a list with the amino acid-specific fragment ions based on the expected M/Z ratio. The peak areas of the TBDMS-derivatized mass fragments were then integrated for the most prominent GC-MS fragments, i.e. (M-15)^+^, (M-57)^+^, (M-85)^+^, (M-159)^+^ and (f302)^+^ using the default settings of MZmine2 (Nanchen et al, 2007). The integrated data was then used to determine the relative abundance of mass isotopomers of individual amino acid fragments using the iMS2flux software tool (Poskar et al, 2012). Error propagation was applied to calculate the mean and standard deviation across 18 measurements for one labelling experiment (two biological times three sample replicates times three GC-MS measurement of each replicate).

Measured mass isotopomer distributions of amino acids were retained, if the following two earlier suggested (Zamboni et al, 2009) quality criteria were met: (i) the ^13^C isotopic abundance in the amino acid carbon backbone in the detected mass isotopomer of an unlabelled biomass sample differs less than 1.5 mol% from the expected isotopic abundance of a carbon backbone with an equal number of atoms; (ii) the fractional ^13^C isotope labelling deviates less than 0.02% from the expected one. To verify the naturally occurring mass isotopomer patterns of the carbon atom backbone in the measured amino acid mass fragments, we quantified the amino acid mass fragments of an unlabelled biomass sample. All amino acid mass fragments, which did not fulfil both evaluation criteria for the unlabelled biomass were not further used. The amino acid fragments such as (M-15)^+^ and (M-57)^+^ report on the same atoms in the amino acid carbon backbone, and thus are indistinguishable to the isotopomer model. Thus, we used only the mass fragment with the highest peak area, assuming a higher accuracy of the isotopic distributions in more abundantly measured mass fragments (Supplementary Data File 5).

### Quantification of intracellular metabolite levels

To determine intracellular metabolite concentrations, 1 mL of cell culture was rapidly quenched by mixing with 12 mL of -40°C pure methanol. The samples were centrifuged (−9°C, 3750 g, 4 min) and the supernatant removed. The cell pellet was frozen in liquid nitrogen and stored at -80°C until extraction. Metabolites were extracted by adding 75% pre-heated ethanol for 3 min at 80°C. During extraction, 200 µl of fully labelled ^13^C-biomass was added as an internal standard (Wu et al, 2005). A vacuum centrifuge (Christ-RVC 2–33 CD plus, Kuehner AG) was used to dry the extracted metabolites for approx. 8h. The dried metabolites were resuspended in 100 µl ddH_2_O. Metabolite measurement was performed according to (Büscher et al, 2009). We used a Waters Acquity UPLC (Waters Corporation, Milford, MA) with a Waters Acquity T3 end-capped reverse-phase column with dimensions 150mm x 2.1mm x 1.8 mm (Waters Corporation). The chromatography was coupled to a Thermo TSQ Quantum Ultra triple quadrupole mass spectrometer (Thermo Fisher Scientific, Waltham, MA) with a heated electrospray ionisation source (Thermo Fisher Scientific) in negative mode with multiple reaction monitoring. Acquisition and peak integration was performed with in-house software and the peak areas were further normalised to fully ^13^C labelled internal standards and the amount of used biomass. The metabolite amounts were normalised to the OD of the cell culture, measured at the sampling point. The normalised metabolite concentrations were converted into intracellular concentration using the dry weight / OD ratio of 0.52 (Sonderegger & Sauer, 2003) and the cell volume / dry weight ratio of 2 mL gCDW^-1^ (Van Urk et al, 1988).

### Fitting of the TSM model to experimental data and variability analysis

The TSM model defined by Eqs. 1-4 and respective variable bounds was fitted to physiological data, metabolite concentrations and Δ_*r*_*G*° values with weighted sum of squares, where the respective measurement errors were used as weights. This fitting was performed as previously (Niebel et al, 2019) with two exceptions: no regularisation was applied and the levels of extracellular metabolite concentrations were allowed to vary between 0 and the extracellular concentration that maximally occurred in the batch experiment (for the metabolites that were consumed and produced). We first estimated Δ_*r*_*G*° values and their respective errors with the Component Contribution Method (CCM) (Noor et al, 2013). However, not all Δ_*r*_*G*° values could be estimated with CCM. Thus, we fitted the TSM model to the physiological and metabolome data, along with the estimated Δ_*r*_*G*° and enforced the first law of thermodynamics to find a complete and thermodynamically consistent set of Δ_*r*_*G*° values. Algebraic equations were used to connect the cell-averaged metabolite concentration measurements to the compartmental metabolic concentrations using known compartment volumes.

After finding a consistent set of Δ_*r*_*G*° and constraining the Δ_*r*_*G*°’s to these values, we used variability analysis to estimate the bounds of all model variables under the experimental condition defined by the fitting. For the variability analysis, we constrained the measured variables (i.e. exchange fluxes and metabolite concentrations) to the values estimated in the fitting plus/minus three experimental standard deviations. For both optimisations problems (fitting and variability analysis), reactions were coupled in groups of reactions that proportionally carried the same flux, as previously done (Burgard et al, 2004). This coupling was done to reduce the computational efforts in the optimisations. Additionally, to efficiently solve the non-linear optimisations, in the fitting and variability analysis, we first solved a linear problem and used that solution as initial guess for the non-linear problem, as done previously (Niebel et al, 2019).

### Atom transition network for isotopomer balancing

^13^C metabolic flux analysis was carried out using the software 13Cflux2 (Weitzel et al, 2013). A 13Cflux2 model with the complete carbon atom transition network was implemented. The metabolic reaction network was the same used for the thermodynamic and stoichiometric model with the exception of the non-carbon compounds and transport reactions. Non-carbon compounds and their respective mass balances were not included in the isotopomer model because ^13^C data traces only carbon atoms. Besides, the different stoichiometric variants of transport reactions were summed to obtain one single reaction for each metabolite transport in the isotopomer model. The atom transitions of the carbon atoms in the 13Cflux2 model were implemented as described (Weitzel et al, 2013) and according to the previously published atom transitions at MetaCYC (Latendresse et al, 2012).

Three instances of the 13Cflux2 models were generated for fitting the measurement data of the [U-^13^C] and [1-^13^C] ^13^C labelling experiments and also their combined data [U+1-^13^C]. In the [U+1-^13^C] ^13^C instance, all reactions were composed of two metabolite variables, mapped either to the carbon flux of the [1-^13^C] or to the [U-^13^C] metabolites. The measured amino acid patterns were associated with their respective metabolite in the cytosol. The isotopomer model and atom transition used are in 13Cflux2 format fml files (Beyß et al, 2019) in Supplementary Data File 7. All reactions in the model were assumed to be fully reversible.

Net fluxes were fixed to the values of the sampled fluxes resulting from the sampling of the flux solution space. Note that because the net fluxes are fixed when we fit the isotopomer model to the ^13^C data, the other excluded balances (i.e. of non-carbon compounds) are still implicitly fulfilled.

### Hit-and-Run sampling

The Hit-and-Run algorithm (Claude et al, 1993) was used throughout our work to sample the flux and concentration polytopes. Here, we describe the steps of this algorithm for the flux polytope with fluxes (*v*) as variables, as it can be generalised for any other convex space. Eq. 1 and the flux bounds define a polytope in *D* = *rank*(*S*) dimensions, where *S* is the stoichiometric matrix. Using the Gauss-Jordan decomposition, we identified *D* vectors of the basis that spanned the polytope. Starting from a feasible flux point *v*_*t*−1_ within the polytope, we extracted a direction *θ* from the uniform distribution on the *D*-dimensional unit sphere and proposed a move *v*_*t*_ = *v*_*t*−1_ + *λ*_0_*θ*. The parameter *λ*_0_ was determined requiring that the proposed flux point satisfied the boundary constraints (flux bounds). First, we needed to compute *λ*_*min*_ and *λ*_*max*_ as the minimum and maximum values of *λ* such that the constraint 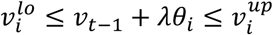 was satisfied for any flux. Then, *λ*_0_ was picked uniformly at random in the interval [*λ*_*min*_, *λ*_*max*_]. This Hit-and-Run algorithm was implemented in MATLAB (R2017a 9.2.0.556344).

As the high-dimensional metabolic space suffers from ill-conditioning due to very different magnitudes along different dimensions, the sampler had difficulty moving along narrower parts of the space. To improve the sampling, we selected directions *θ* from the axes of an approximate Lower-John (De Martino et al, 2015) ellipsoid containing the polytope instead of the unit sphere. The equation of the ellipsoid was obtained from Expectation Propagation algorithm (see below).

### Division of the flux solution space into sectors

The division of the solution space of fluxes, concentrations, and Δ_*r*_*G* described by stoichiometry and thermodynamic constraints (TSM model) into two polytopes recreated the occurrence of thermodynamically infeasible loops (Beard et al, 2002) in the flux solutions sampled from the flux polytope. It has been demonstrated by using known theorems of the alternatives (Schrijver, 1986) that the existence and computability of Δ_*r*_*G* is possible if and only if reaction flux configurations are devoid of closed loops (Beard et al, 2002;De Martino, 2013;Noor et al, 2012;Price, N. D. et al, 2002). To avoid such loops, we applied the Monte Carlo method described in (De Martino et al, 2013) to detect and remove such infeasible flux sign configurations beforehand, or at least the ones that affected the most the acceptance rate of the sampling. Implemented in C++, this method sampled 100 uncorrelated points per second and found infeasible loops in 0.5 ms with an Intel quad-core and clock speed of 3.8 GHz. In the model used here, we detected four major loops (Supplementary Table 1) whose removal lead to sampling feasible configurations with workable acceptance rates. Loops were removed by considering for the sampling only the flux sign configurations that excluded them, leading to the exclusion of specific orthants of the flux solution space. Through manipulation of the flux bounds of nine reactions, we generated 16 disjoint polytopes whose combination constituted the complete feasible flux solution space. The so-called sectors (Supplementary Table 2) were then almost devoid of loops and we could sample them independently. We checked that further loop removal did not lead to consistently increased acceptance rates for the sampling, while on the other hand it would eventually increase exponentially the number of sectors to sample.

### Determination of volume and ellipsoid of the flux solution space through Expectation Propagation

We applied Expectation Propagation (EP) to calculate the volume of individual sectors in the flux solution space, and to derive approximating ellipsoids that were used to scale the flux and concentration polytopes during Monte Carlo sampling. Expectation Propagation (EP) (Minka, 2001) is a powerful and computationally efficient algorithm that approximates an intractable, high-dimensional target distribution by a tractable one. Typically, the approximating distribution is a multivariate Gaussian with mean *μ* and covariance matrix *C* (denoted by *N*(*μ, C*)), in which case EP is used to obtain the values of *μ* and *C*. Recently it was shown that EP provides an accurate multivariate Gaussian approximation to the uniform distribution on the convex polytope defined by the solution set of an underdetermined linear system *Ax* = *b*, where *x* is allowed to vary within a bounded set (Braunstein et al, 2017).

Based on the results of (Braunstein et al, 2017) we used the “zero-temperature” implementation of the EP scheme (Braunstein A et al, 2019) to provide a multivariate Gaussian approximation, *N*(*μ*_*v*_, *C*_*v*_), to the solution space of each flux sector. To derive a proxy for the sector volume, we used the exponential of the Shannon entropy *H* of the multivariate Gaussian distribution, given by 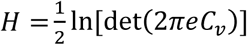 (Cover & Thomas, 2006). The volumes of the different sectors of WT and TM6 flux spaces are provided in Supplementary Table 2.

We further used EP to obtain multivariate Gaussian approximations for each of the flux sectors and the concentration polytope defined for a fixed flux configuration. The covariance matrix of each Gaussian approximation defines an ellipsoid that matches the underlying polytope (De Martino et al, 2015). This matrix was then used to rescale the corresponding polytope in order to increase the speed of Monte Carlo sampling (De Martino et al, 2015).

### Parallel tempering

Parallel Tempering (PT) is an MCMC-based algorithm that enables the sampling of target distributions *p*(*x*) which are difficult to sample with just a single Markov chain (Swendsen & Wang, 1986). The general idea of PT is to generate *n* parallel Markov chains (“replicas”), where *n* − 1 chains sample distributions that are more diffuse than the distribution of interest, *p*(*x*). These distributions are obtained by varying a “temperature” parameter *T*, with *T* = 1 corresponding to the distribution of interest and *T* > 1 resulting in a distribution more diffuse than *p*(*x*). Chains corresponding to higher temperatures are able to explore larger volumes of the space compared to chains of lower temperature. By allowing the chains at different temperatures to exchange points while preserving their joint distribution, the lower-temperature chains are thus able to access points of the space that would be difficult to reach with just the lowest-temperature chain alone.

In this work, our target was the uniform distribution over each sector of the flux polytope (Polytope A) conditional on the concentration polytope (Polytope B), a set denoted by A∩B. Normally, the sampling of Polytope A sectors can be efficiently accomplished by a Hit-and-Run MCMC sampler after proper rescaling of the flux space (De Martino et al, 2015; Fleming et al, 2012), as described above. However, the requirement to sample flux configurations in Polytope A which also result in a non-empty Polytope B (a feasible set of thermodynamic constraints defined by Eqs. 2-4 and their respective variable bounds), makes the sampling considerably more difficult, and a simple rejection sampling approach becomes computationally infeasible. We therefore resorted to a PT-based strategy described in detail below.

For a given flux point *v*, the thermodynamic constrains (Eqs 2-4) form a polytope that can be described as the set of *x* ≡ {Δ_*r*_*G, lnc*} which satisfy the equality constraint *Ax* − *b* = 0 subject to bounds [*x*^*lo*^, *x*^*up*^]. The matrix *A* can be derived from Eqs. 3 and 4, while the bounds can be obtained from the bounds on Δ_*r*_*G* and *lnc*, together with the bounds implied by Eq. 2. For the construction of the diffuse target distributions of our PT scheme, we first introduced a vector *ε* which quantifies the violation of each of the equality constraints. For a given flux point, the minimal sum of absolute constraint violations can then be calculated by solving

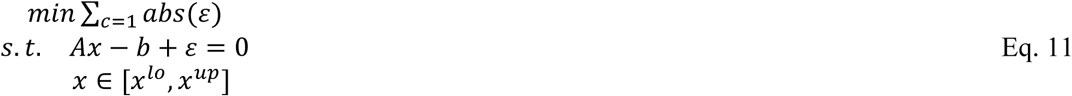

where ‖*ε*‖_1_ denotes the L1 norm of *ε*. The minimum of ‖*ε*‖_1_ quantifies how close a given flux point is to satisfying the equality constraints. Flux points belonging to *S*_*A*|*B*_ (the target set) correspond to ‖*ε*‖_1_ = 0. Moreover, flux points with small ‖*ε*‖_1_ lie closer to *S*_*A*|*B*_ than fluxes with large ‖*ε*‖_1_.

For our PT scheme, we defined a set of parallel Markov chains in the flux space (Polytope A) for a set of different temperatures. Each chain sampled flux points whose minimal violations *ε* (computed from Eq. 11) are distributed according to a zero-mean multivariate normal distribution with independent components. The *i*-th component of this distribution corresponds to the *i*-th equality constraint, and has marginal distribution of the form 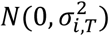, i.e. the normal distribution with mean 0 and variance 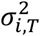 depending on the temperature parameter *T*. More specifically, the variance of each component was set equal to the squared temperature parameter times a very small target variance 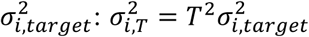. In this way, points sampled by the replica at temperature *T* = 1 lie very close to *S*_*A*|*B*_ and can thus be thought to correspond to the target distribution. On the other hand, points sampled by replicas at *T* > 1 correspond to greater constraint relaxations and therefore lie in a set larger than *S*_*A*|*B*_, which can be much easier to explore than *S*_*A*|*B*_ itself.

To define 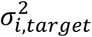 for the *i*-th error component, we first sampled 10,000 flux points from Polytope A without the constraints of Polytope B, and calculated the minimal violation vector *ε* for each of these points by solving the optimisation problem Eq. 11. For each component *ε*_*i*_ of *ε*, we located the 10% and 90% percentiles of the resulting distribution and selected the percentile with the largest absolute value, denoted by *M*_*i*_. We finally set 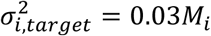. To obtain the higher-temperature replicas, we chose different values of *T* above 1 and below 900, resulting in a set of parallel replicas in the flux space. Depending on the flux space sector, six or seven replicas were used for PT sampling.

To implement the PT scheme, two types of updates are necessary: one to determine the movement of a replica at a given temperature and another for swapping the states of replicas at different temperatures. At each temperature *T* and step *k*, a move from *v*_*k*_ to *v*_*k*+1_ was proposed by a Hit-and-Run algorithm as described below, and accepted with probability

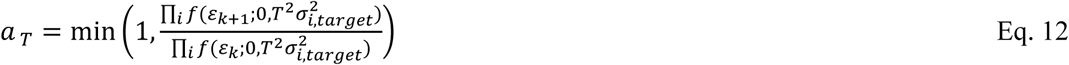

based on the minimal violation vectors *ε*_*k*+1_ and *ε*_*k*_ (computed from Eq. 11 for *v*_*k*+1_ and *v*_*k*_). In the above equation, *f*(*x; μ, σ*^2^) denotes the probability density function of the normal distribution with mean *μ* and variance *σ*^2^.

The swap of flux points *v*_*k*_ and 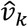 at step *k* between two replicas at different temperatures, *T* and 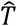, was accepted with probability

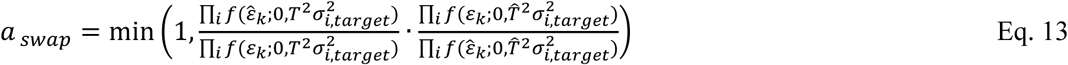

based on the minimal violation vectors *ε*_*k*_ and 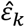 (computed from Eq. 11 for *v*_*k*_ and 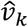).

The six temperature levels were manually adjusted so that the swap rates (fraction of accepted swaps between temperatures over total number of attempts) were similar among pairs of neighboring temperatures. Each temperature level could only swap flux points with the level immediately above or below. High swap rates between pairs of temperatures make the two temperature levels redundant, while low swap rates prevent the efficient flow of information between temperature levels. We therefore aimed for swap rates of around 20% between all pairs of temperatures. The manual adjustment of the temperature levels was carried out for each sector of the flux space.

The overall movement of the PT sampler was configured as follows: the replica of each temperature was allowed to perform 5 Metropolis-Hastings (M-H) updates (with acceptance probability given by Eq. 12), after which a pair of temperatures was chosen uniformly at random and a swap was proposed (with acceptance probability given by Eq. 13). Each M-H update at fixed temperature consisted of a number of Hit-and-Run moves which allowed the corresponding chain to move a sufficient distance away from the current point before attempting to accept a new point, resulting in better exploration of the flux space. The number of Hit-and-run “skips” was set to 3, 10, 100, or 1000 depending on the temperature.

PT sampling was performed in MATLAB (R2017a 9.2.0.556344) running on two different computer clusters: a local cluster with 48 cores (clock speeds between 2 and 3.3 GHz; 4/8 GB of RAM per core) and the Peregrine HPC cluster of the University of Groningen (6 cores with clock speeds between 2.2 and 2.6 GHz and 128 GB of RAM per core). The sampling of the different temperatures was done in parallel in different cores using the Parallel Computing Toolbox of MATLAB. With these settings, the sampling of one sector took on average 3.5 hours, with 30000 swaps performed among temperatures to generate 30000×5 points for the lowest temperature.

### UMAP

To visualise the sampled flux solution space, we applied Uniform Manifold Approximation and Projection (UMAP) (McInnes et al, 2018), which is a dimension reduction algorithm that represents multi-dimensional spaces in fewer dimensions while keeping a similar distance between points as the original multi-dimensional space. Results in Fig. 4c were generated through UMAP in python 3 with 50 neighbours and minimum distance of 0.5.

### Statistical analysis

To identify differences in flux distributions between two sets of samples, we performed different statistical tests. For the comparison of the complete flux sample (1000 points obtained from sampling the flux solution space) to a sub-set of 200 net fluxes samples best-fitting to ^13^C-data, we generated 10000 random sub-samples of 200 net flux samples from the complete flux sample and we calculated the medians and IQRs of each flux in each sub-sample. If the median of a flux in the sampled subset of best-fitting net flux samples was within the 99.99% percentile of the distribution of medians from the random sub-samples, then the flux median was considered as not significantly different between the complete sample and the sub-sample. The same procedure was applied to the interquartile range.

To compare the flux distributions obtained from best fitting net fluxes to the ^13^C-data with and without the flux-force constraint, we used a Kolmogorov-Smirnov statistic (KS) to qualitatively measure the difference between two sub-samples. To this end, we generated pairs of random sub-samples of 200 points from the complete flux sample. The KS statistic was calculated for each pair of random sub-samples. Similarly to the previous approach, if the KS statistic calculated from the two sub-samples best fitting ^13^C-data with and without flux force constraint was inside (99.99% percentile) the distribution of random KS statistics for a particular flux (Fig. 6d), such flux was considered as not significantly different between the two sub-samples. All sub-samples mentioned were chosen based on ^13^C-data fitting with the size of 20% of the complete flux sample (i.e. 200 points) obtained from the flux solution space. All statistical analyses were performed with python 3.

### Optimisations with 13Cflux2

The fits of the isotopomer model to ^13^C data were performed with 13Cflux2 (Weitzel et al, 2013). Besides when stated otherwise in the text, the net fluxes from sampling were fixed in the optimisation leaving the labelling exchange fluxes as free variables. No labelling exchange fluxes were excluded (set to zero) from the model. Labelling exchange fluxes were bounded between 0 and 1000, besides when flux-force constraint was applied, where the lower bounds changed in a reaction-basis according to the value of sampled Δ_*r*_*G*.

As it is common practice, each optimisation run with 13Cflux2 was performed several times with different starting points to increase the chance of finding a global optimum. Besides the number of starting points, another parameter we needed to tune was the number of maximum iterations for the optimisation problem. In theory, we wanted both the number of starting points and the number of maximum iterations to be as high as possible, however, these optimisations are complex and very time consuming. Having more starting points implies more optimisation runs and larger number of iterations implies longer runs. To find the ideal combination we evaluated the optimisation residual (objective function) and the total optimisation time for each net flux. We tested three combinations of starting points/ maximum iterations, respectively: 200/50, 100/100, 50/200. All tests were made for 10 net flux samples fitted to the [U-^13^C] data set. We found that 200 iterations and 50 starting points yielded the lowest optimisation residuals in acceptable amounts of time (Fig. S4a,b). The starting points were obtained from sampling the labelling exchange flux space for each net flux point, independent of the labelling measurements, with an HR algorithm (13Cflux2 *ssampler* function). The fit was performed with the function *fitfluxes* of 13Cflux2.

Optimisation times were, in average, 15 minutes per fit to [U-^13^C] or [1-^13^C] and 26 minutes to [U+1-^13^C]. All optimisations were run on a computational cluster with 48 cores and clock speeds between 2 and 3.3 GHz. The sampling of the starting points is extremely fast and thus its running time is negligible compared to the total optimisation time.

## ACKNOWLEDGMENT

MH acknowledges funding from the European Union Seventh Framework Programme (FP7-KBBE-2013-7-single-stage) under grant agreement no. 613745 (PROMYS), from the European Union H2020 Programme under grant agreement no. 675585 (SymBioSys), and the Dutch Research Council (NWO) for the Systems Biology Center for Energy Metabolism and Aging. DDM acknowledges a short-term EMBO fellowship (no. 7414). Gesa Behrend’s support in the consolidation of the 13Cflux2 model is kindly acknowledged.

## CODE AVAILABILITY

The computational code and scripts are available at https://github.com/molecular-systems-biology.

## SUPPLEMENTARY FIGURES

**Figure S1:**
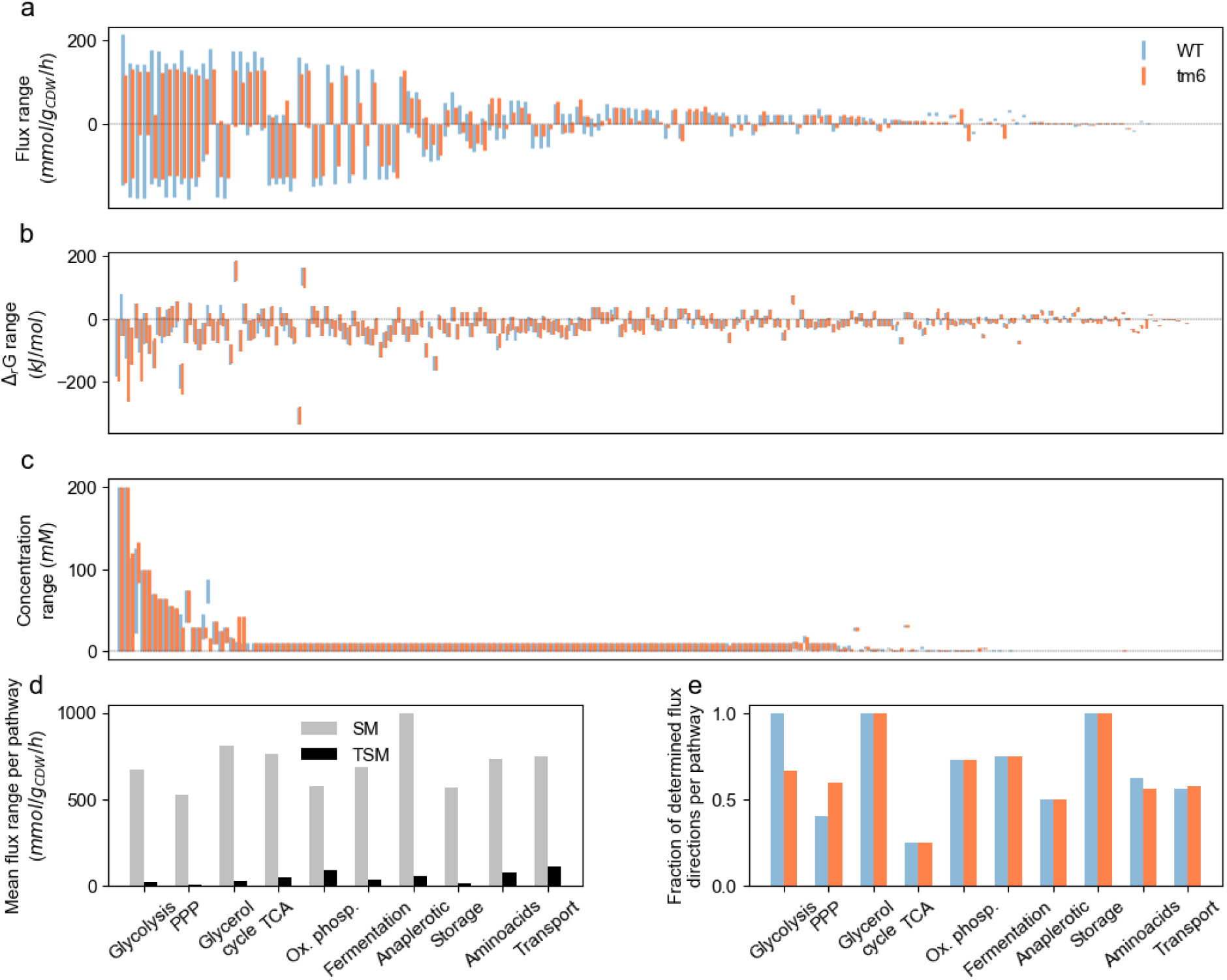
Ranges of variables and flux directions determined by variability analyses. (a), (b), (c) Flux (a), Gibbs energy of reaction (b) and concentration (c) ranges determined from the variable bounds determined by variability analyses for WT (in blue) and TM6 (in orange); in (b) the range of reaction ergstSYN is omitted for visualisation purposes (−1044 to - 927 KJ/mol), and in (c) the concentration of NAD in the cytosol (10 to 1000 mM). x-axes in (a), (b) and (c) sorted by decreasing WT range. (d) Mean flux range per pathway determined from the variability analysis results for WT with a stoichiometric model (SM, in grey) and physiology data, and additionally thermodynamics and metabolome data (TSM, in black). (e) Fraction of determined flux directions per pathway, determined from the variability analysis with the TSM model, for WT (in blue) and TM6 (in orange).

**Figure S2:**
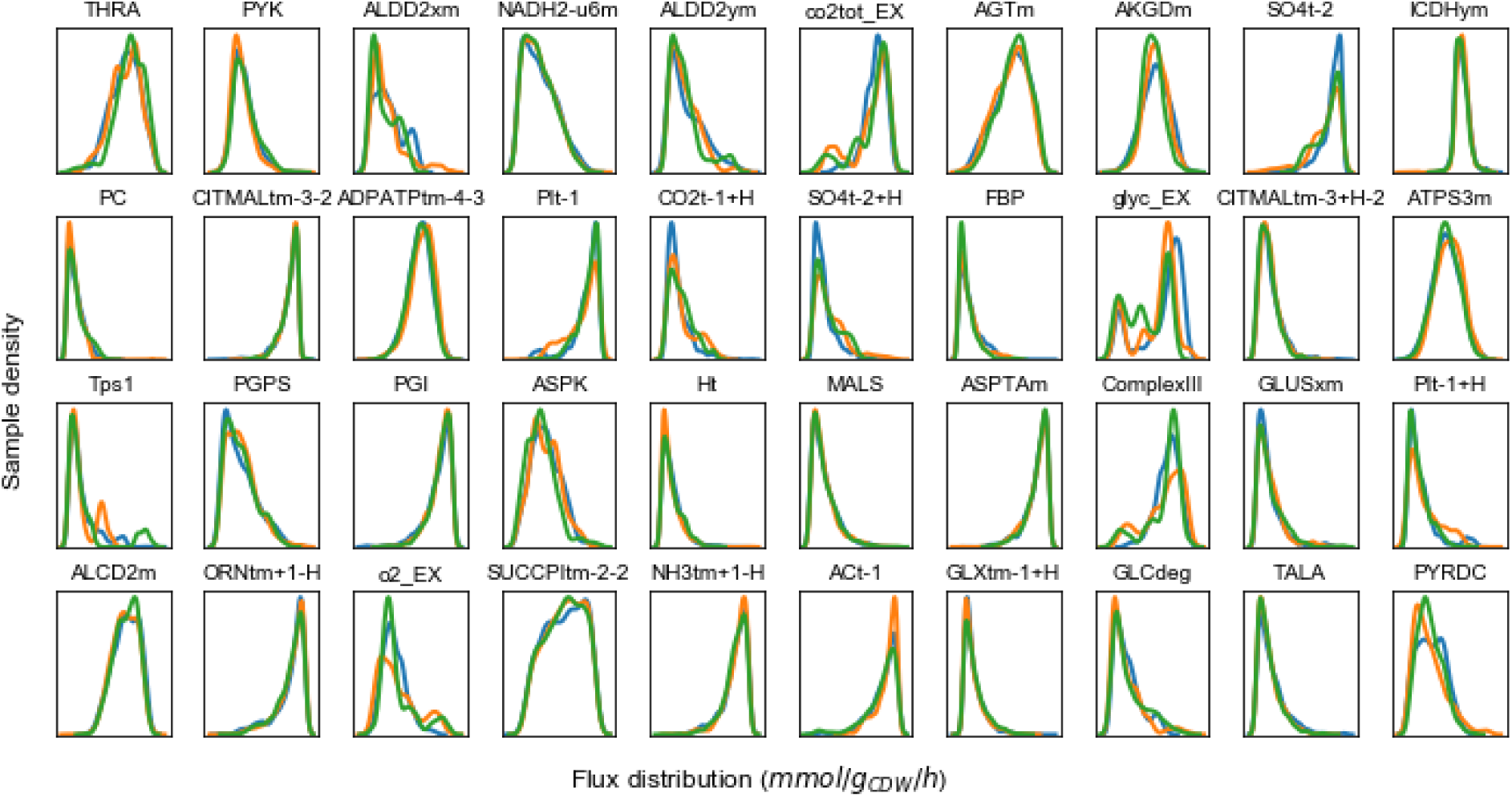
Sampler reproducibility. Sample density of 40 random flux distributions for three independent sampling runs with the sampling method devised in this work, using Hit-and-Run algorithm and Parallel Tempering (in three colours: blue, orange, and green) for TM6 flux solution space. Names of reactions are in Supplementary Data File 1.

**Figure S3:**
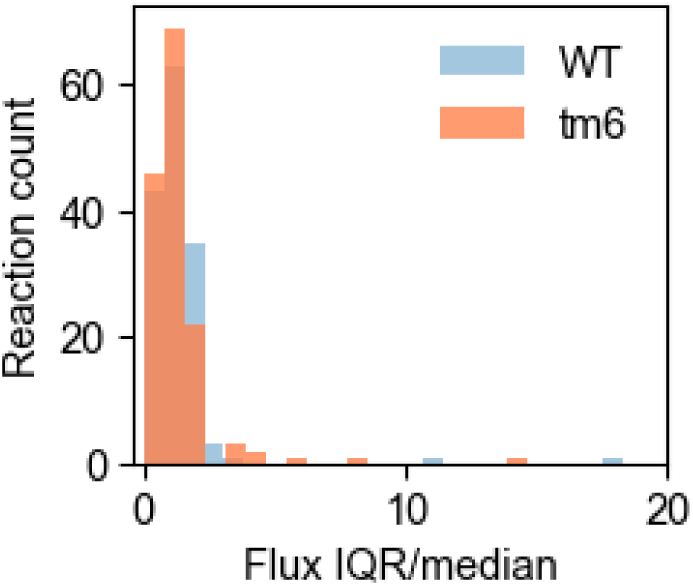
IQR/median of sampled fluxes for WT and TM6. Interquartile range (IQR) over median of sampled fluxes for WT (in blue) and TM6 (in orange). Sampled fluxes obtained from sampling the flux solution space conditional on the concentration space with Hit-and-Run and Parallel Tempering. Here, only IQR/median values up to 20 are shown, leaving out 1 and 3 reactions for WT and TM6 respectively.

**Figure S4:**
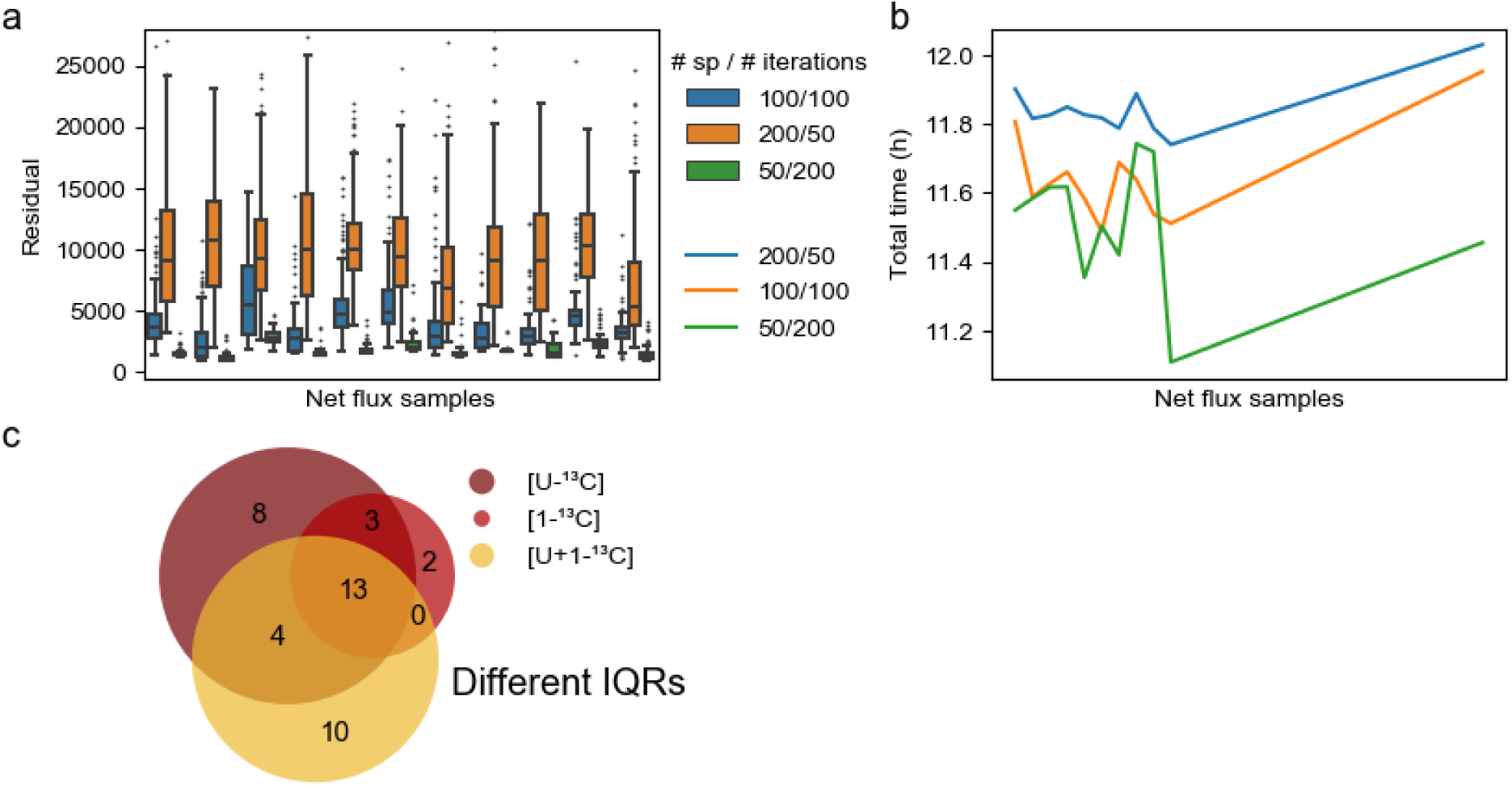
Tuning and effect of optimisations to fit isotpomer model to ^13^C labelling data. (a) Distribution of optimisation residuals with number of starting points (n_SP) and maximum number of iterations (n_iter) for different net flux samples. Residual value is normalised to the overall minimum value for all optimisations. (b) Total optimisation time (y-axis) for different sampled net flux points (x-axis) with various combinations of n_SP and n_iter. Total time corresponds to the sum of optimisation time for all starting points in each n_SP spec. All optimisations were run on a computational cluster with 48 cores and clock speeds between 2 and 3.3 GHz. (c) Number of reactions for which the flux distribution IQR was statistically different, when comparing before and after scoring, for the different experiments. Scoring represents the selection of the 20% best fitting net flux samples, which are then statistically compared to the full sample. A number of random distributions, of the same size of the 20% best fitting sample, are generated from the complete net flux sample. From these random distributions, interquartile ranges (IQR) are calculated for each reaction flux. Reactions for which the IQR of the 20% best fitting sample is outside (99.99%) the distribution of IQRs from the random distributions, is considered to present a significant statistical difference. Reactions identified in overlapping circles represent the same reactions identified in the two or three labelling experiments.

**Figure S5:**
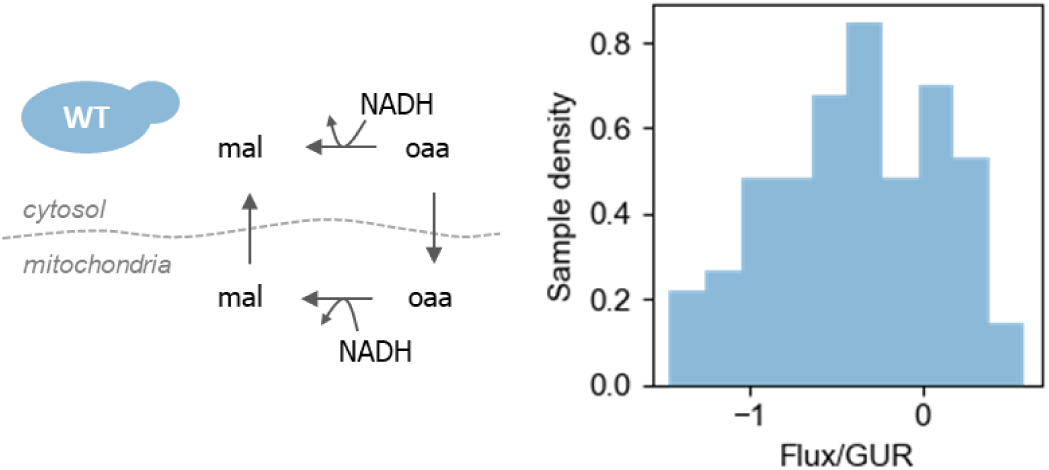
Uncertainty in flux estimation for malate-oxaloacetate shuttle in wildtype. This shuttle would be present in WT if the cytosolic malate dehydrogenase reaction (top-most arrow) would go from malate to oxaloacetate. The median of the flux goes in the opposite direction, however its flux distribution crosses the 0 making it uncertain if this shuttle is present in WT. Flux distribution from net flux samples scored with the 20% best fits of the isotopomer model to [U+1-^13^C] with flux force constraint. Flux values are normalised with glucose uptake rate (GUR). mal: malate, oaa: oxaloacetate.

**Figure S6:**
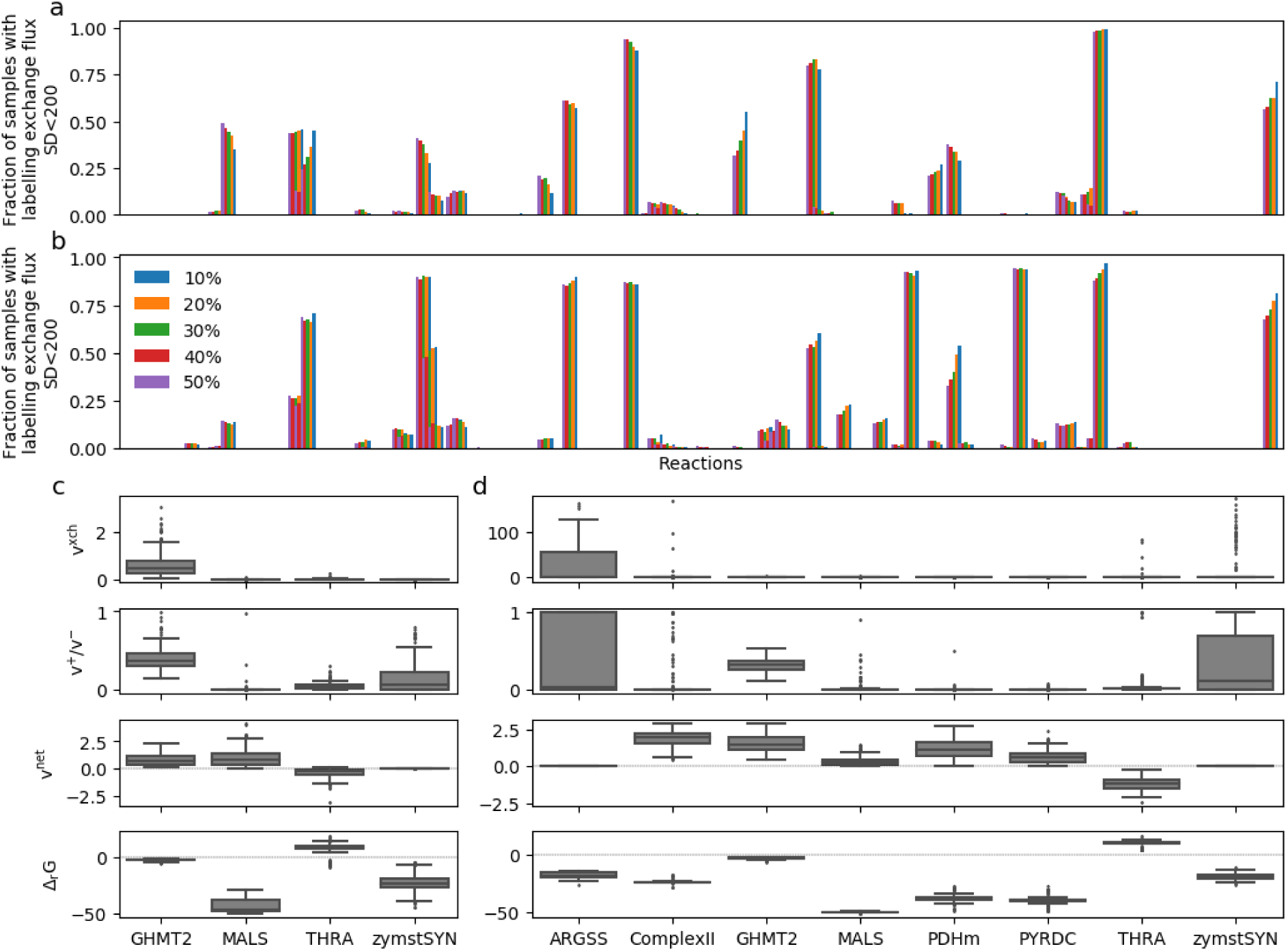
Identification of labelling exchange fluxes. (a), (b) Labelling exchange fluxes were considered determined for WT (a) and TM6 (b) if at least 50% of the labelling exchange flux (y-axis) solutions (^13^C-optimisations of 20% best scoring fits) were below the empirical threshold of 200 mmol/g_CDW_/h. This was tested for different selections of the best residuals according to the fitting of the isotopomer model to [U+1-^13^C] data and using the flux force constraint, from 10 to 50% of the net flux samples. (c), (d) Distributions of labelling exchange fluxes (v^xch^), ratio of forward and backward fluxes (v^+^/v^-^), net fluxes (v^net^) and Gibbs energies (Δ_*r*_*G*) for the reactions with considered determined labelling exchange fluxes for WT (c) and TM6 (d). Distributions show only the 20% best samples according to the residual of fitting the isotopomer model to [U+1-^13^C] data and using the flux force constraint. Δ_*r*_*G* values used to constrain the labelling exchange fluxes in the fitting.

**Figure S7:**
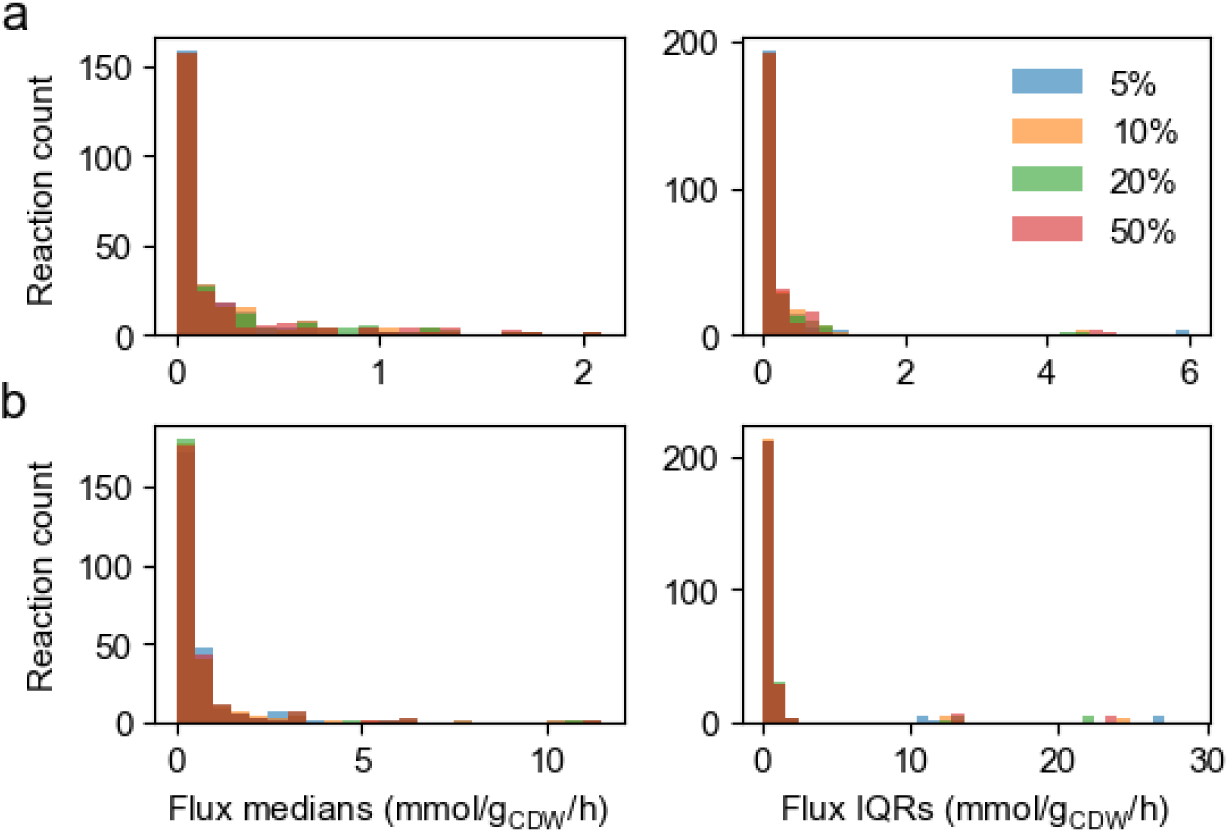
Flux statistics at different scorings with ^13^C data. Flux medians (left plots) and interquartile ranges (IQRs, right plots) in WT (a) and TM6 (b) at different scorings of the net flux samples with the fitting to the [U+1-^13^C] data with flux force constraint.

## Supplementary information – Gibbs energy balance in thermodynamic and stoichiometric model

Once the standard Gibbs reaction energies are calculated and considered as fixed in the model, the Gibbs energy balance (Eq. 4) is automatically fulfilled through the other constraints in the TSM problem. Eq. 4 states that the intracellular energy production equals the energy exchange rate with the exterior of the system. The total intracellular Gibbs energy dissipation is defined as the sum of the product of intracellular fluxes with the respective Gibbs energy of reaction, while the Gibbs energy exchange rate is defined as the same but for exchange reactions. This is actually a consequence of mass balance and thermodynamic definitions as demonstrated in equations S1 to S4.

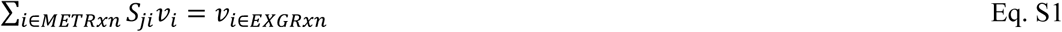

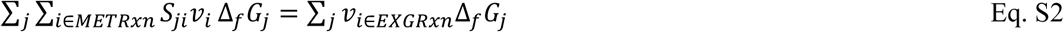

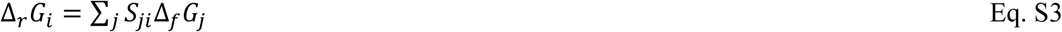

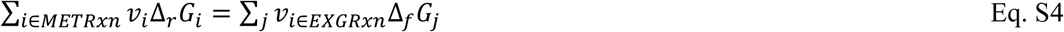

*S, v*, Δ_*r*_*G* and Δ_*f*_*G* are the stoichiometric matrix, metabolic flux, Gibbs energy of reaction and Gibbs energy of formation, respectively. Indexes *i* and *j* represent reactions and metabolites. Here, reactions are divided in 2 sets: *METRxn* and *EXGRxn* represent intracellular and exchange reaction sets respectively. Eq. S1 is a reformulation of Eq. 1 where exchange reactions are moved to the right side of the equation. Exchange reactions are defined as having one single substrate and no product, thus *S* = −1 and there is no sum over reactions in Eq. S1 because there is one single exchange reaction for one metabolite. Eq. S2 is an extended version of Eq. S1 to which the sum over metabolites and multiplication (inside the sum) with Gibbs formation energy were implemented on each side of the equation. Eq. S3 states that Gibbs energy of reaction is the sum of the Gibbs formation energies of all the metabolites involved in that reaction. Using Eq. S3 in Eq. S2 we generate Eq. S4 which is the Gibbs energy balance. This demonstration proves that we do not need to enforce Eq. 4 in our TSM constraints. This is true once we fix Δ_*r*_*G*^0^ values to the result of the first fitting.

This demonstration proves that, in theory, the Gibbs energy balance should be naturally fulfilled even when not explicitly formulated, for a fixed Δ_*r*_*G*^0^ set. However, when we do not enforce it, we get a negligible that, even though small, is not zero. This happens because in the first fitting we find Δ_*r*_*G*° values “around” their value from the component contribution method considering a measurement error, but we consider Δ_*f*_*G*^0^ as a fixed value. After this fitting, we fix Δ_*r*_*G*^0^ values that are slightly different then their measurement mean, which causes a small disparity between the values of Δ_*r*_*G*^0^ and Δ_*f*_*G*^0^.

## Supplementary tables

**Supplementary Table 1.** Most frequent loops in the metabolic network within flux bounds calculated from variability analysis for the experimental data in this work (both WT and TM6). Reactions involved in each loop and corresponding sign pattern causing thermodynamic infeasibility. + and – represent a positive and negative flux, respectively, relative to direction defined in the model. As an example of how to read the table: the last row means that if the flux of Htm and PYRtm1 are positive and the flux of PYRtm2 is negative, any flux solution is thermodynamically infeasible. ACONT: cytosolic aconitate hydratase, ACONTm: mitochondrial aconitate hydratase, CITICITtm: combined transport of citrate and isocitrate between cytosol and mitochondria, Htm: hydrogen transport between cytosol and mitochondria, CITMALtm1/2: two different variants of combined transport of citrate and malate between cytosol and mitochondria, ACtm1/2: two different variants of acetate transport between cytosol and mitochondria, PYRtm1/2: two different variants of pyruvate transport between cytosol and mitochondria.

| Reactions | Sign pattern |
| --- | --- |
| ACONT<br>ACONTm<br>CITICITtm | - |
|  | + |
|  | + |
|  | + |
|  | - |
|  | - |
| Htm<br>CITMALtm1<br>CITMALtm2 | + |
|  | + |
|  | - |
| Htm<br>ACtm1<br>ACtm2 | + |
|  | + |
|  | - |
| Htm<br>PYRtm1<br>PYRtm2 | + |
|  | + |
|  | - |

**Supplementary Table 2.**
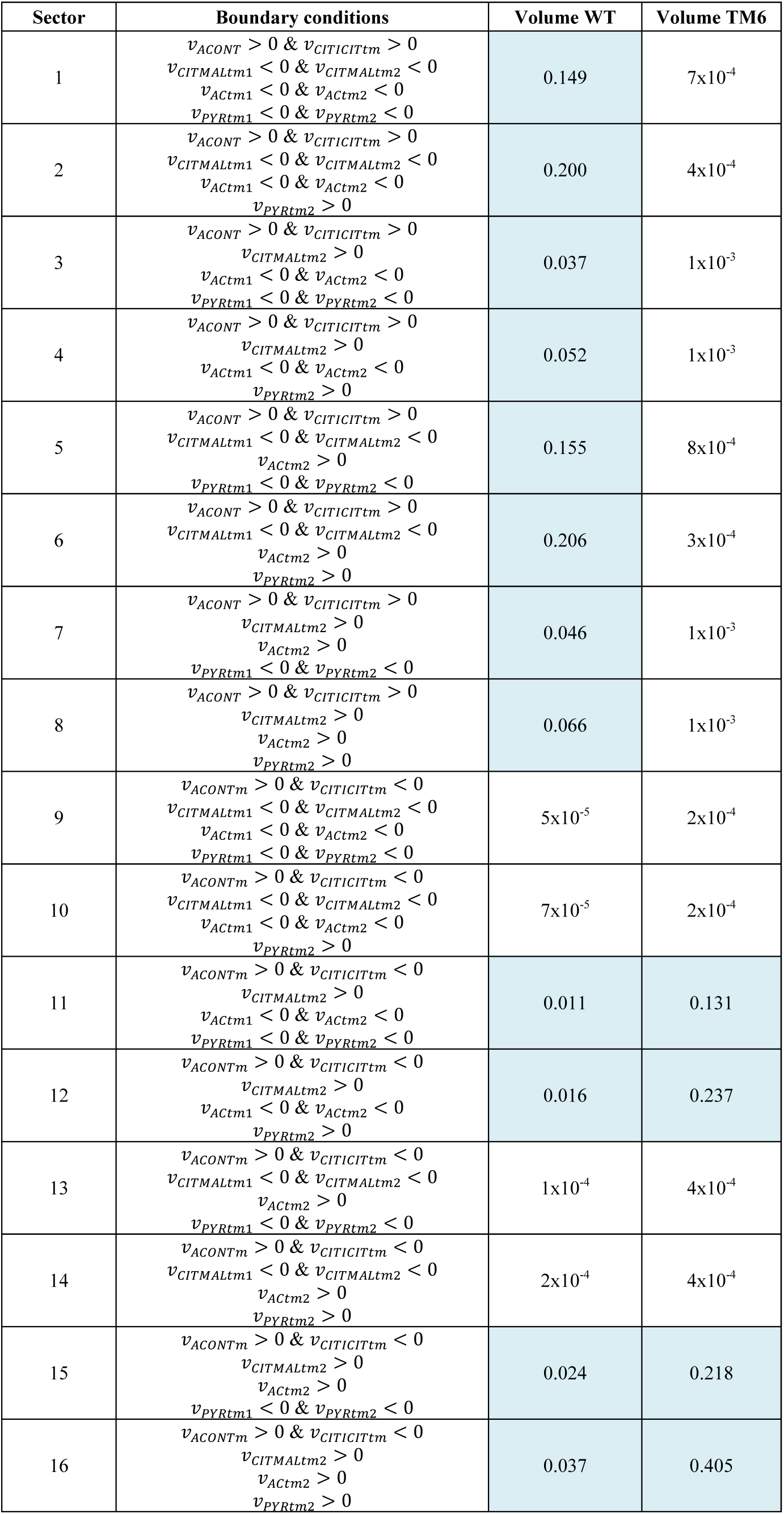
Volume of the sectors of the flux solution spaces of WT and TM6. This volume is calculated through Expectation Propagation, and it is a relative measure used to compare the size of different solution spaces but with no physical meaning on its own. Boundary conditions define flux boundaries of the different sectors. For sampling, only sectors with volume above 0.01 (blue shading) were considered. The rest of the sectors would likely not contribute to the final sample of the flux solution space, and were thus discarded. For WT, the 12 largest sectors covered 99% of the total volume, while in TM6 the 4 largest sectors covered 99%.

